# Charting Taxa in *Amanita* Section *Amidella* (*Basidiomycota : Amanitaceae*)

**DOI:** 10.1101/2024.07.30.605754

**Authors:** Paulo Oliveira, Ricardo Arraiano-Castilho

## Abstract

Species determination in the *Amidella* clade is notoriously difficult, because of the relative dearth of diagnostic characters and the rather common occurrence of homoplasies. This results in a substantial number of misnamed and unnamed collections, a misapprehension of the geographic range of known species, and a gross underestimation of the number of species it contains. To assess the diversity that should be considered as part of *Amidella*, DNA sequences available for this group were retrieved in public nucleotide databases, using a combination of approaches to achieve a comprehensive representation. Phylogenetic analysis based on the aligned ITS sequences, consistently with the results from other molecular markers (ncLSU, *RPB2*, *TEF1*, *BTUB*), suggests five major clades: one containing the type species *Amanita volvata*, another for *Amanita ponderosa* and allies, a third one (roughly half of all species) with *Amanita lepiotoides*, and two others without valid species yet. At species level, around 81 clades were delimited, of which only 16–17 can be assigned a valid name, with a few more corresponding to provisional taxa listed in the amanitaceae.org website. Up to three further species without assigned sequences might correspond to the proposed clades. The current evidence suggests a rather narrow geographic range for most of these clades. This study provides a phylogeny-arranged outlook of the worldwide distribution of *Amidella* species, and an infrasectional framework for optimizing taxonomic sampling and designing clade-specific molecular markers to assist in identification.

**Graphical abstract:** 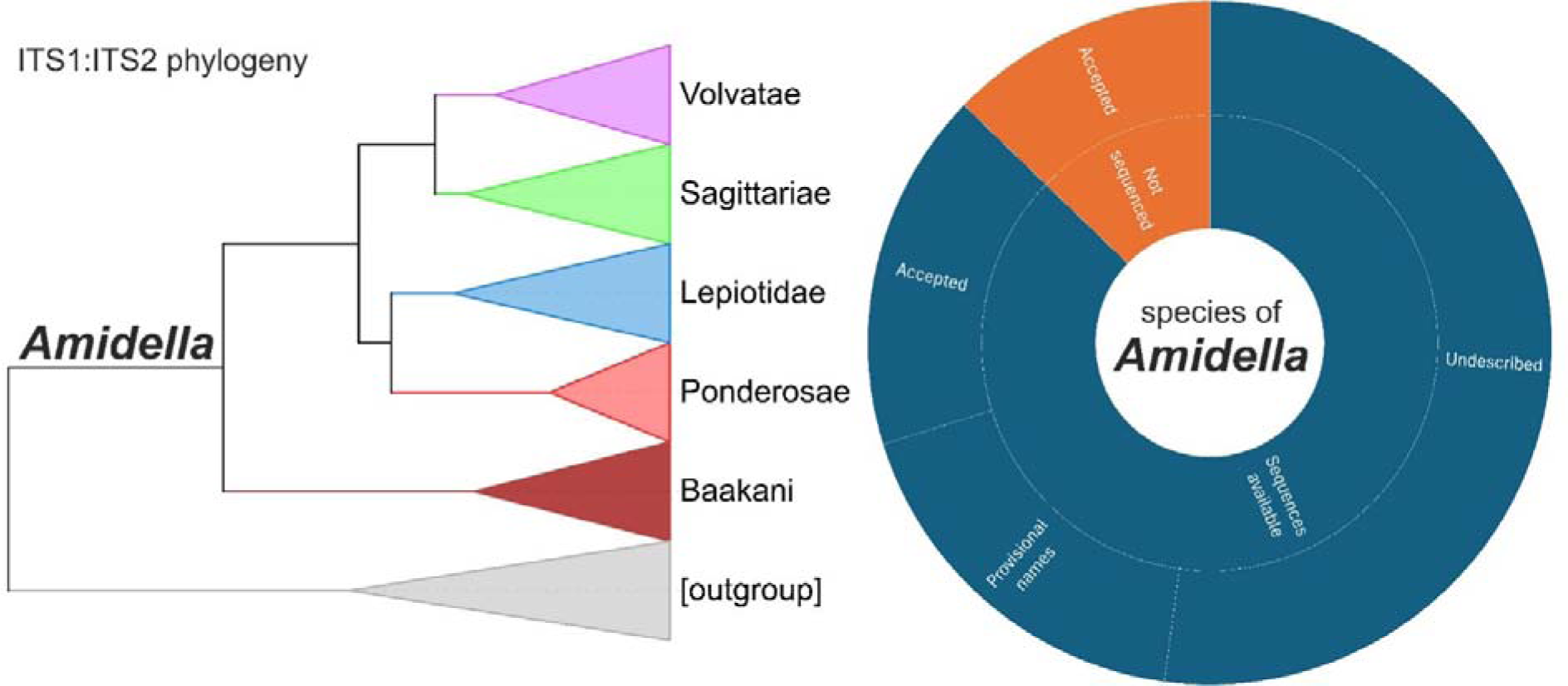

## Introduction

The creation of *Amidella* as a taxonomic group dates from Edouard-Jean Gilbert’s initial proposal as a separate genus, within family *Amanitaceae* created by the same author, and subsequently as a subgenus in *Amanita* Pers. (Neville & Poumarat 2004). The name underscores the importance attached to the basidiospore amyloid reaction with iodine which the name refers to (Konrad and Maublanc 1948).

The current taxonomic arrangement within *Amanita* recognises *Amidella* at the section level in subgenus *Amanitina*, bracketed by sections *Roanokenses* and *Arenariae* (Cui et al. 2018). This is the culmination of a process whereby several groups with amyloid basidiospores segregated from the original concept of *Amidella* (Wolfe et al. 2012; Cui et al. 2018; Riccioni et al. 2019), leaving the name *Amidella* solely for a morphologically distinctive, relatively small group of species (Cui et al. 2018; Tulloss and Yang 2024).

Its demarcation from other congeneric groups is such that identification in the field as ‘*Amidella’* can be straightforward, but in contrast reaching a determination at species level is very challenging (Tulloss and Yang 2024). It may be achieved at regional level, as done recently for Europe with prominent recourse to non-morphological characteristics such as soil, vegetation and phenology (Arraiano-Castilho et al. 2022). This notwithstanding, too many specimens have been assigned equivocally with species epithets such as *volvata*, *avellaneosquamosa* or *curtipes*. The problems arising from this are very difficult to tackle, and the only compelling argument against misidentifications have been the evident phylogenetic disparities (Moreno et al. 2008; Kim et al. 2013; Cho et al. 2015; Cui et al. 2018).

The ingrained notion of *Amidella* as a small group of species, in spite of some evidence against it (Cui et al. 2018; Tulloss and Yang 2024), favours the continued misapplication of known epithets and does not encourage the exploration of any meaningful taxonomic subdivisions. Partial phylogenetic sampling, not to mention mere blast searches, does little to address these issues; a comprehensive overview of the diversity of *Amidella* is definitely needed.

This study attempts such comprehensiveness based on publicly available molecular databases. The approach cannot cover a few known species lacking molecular representation, but it provides a novel picture of the group, establishing its boundaries, recognising more than 80 clades at species level, most of them undescribed, and demonstrating a phylogenetic substructure of great value for separating sympatric species. Knowledge gaps, such as the need for new morphological criteria and the underrepresentation of vast geographical regions, are clearly defined under the new taxonomic view provided here.

## Materials and Methods

### Adopted identifications

There are concerns about the correct application of some names. Following known authorities of genus *Amanita*, a tentative set of sequenced specimens are adopted as reference (Table 1).

**Table 1.** Outline of the nucleotide accessions representing names that are frequently misapplied.

| Current name (basionym) | GenBank accessions | Authority |
| --- | --- | --- |
| *Amanita volvata*<br>(*Agaricus volvatus* Peck 1872) | AF024485 | Weiß et al. (1998)<br>Tulloss, R. E.<br>http://www.amanitaceae.org/?Amanita%20volvata |
| *Amanita peckiana*<br>(*Amanita peckiana* Kauffman 1913) | HQ539720 | Tulloss, R. E.<br>http://www.amanitaceae.org/?Amanita%20peckiana |
| *Amanita curtipes*<br>(*Amanita baccata* f. *minor* Bres. 1928) | EF653963<br>EF653960 | Moreno et al. (2008) |
| *Amanita avellaneosquamosa*<br>(*Amanitopsis avellaneosquamosa* S. Imai 1933)<sup>a</sup> | MH508259<br>AF024441<br>MH486380<br>MH508682 | Weiß et al. (1998)<br>Tulloss, R. E.<br>http://www.amanitaceae.org/?Amanita%20avellaneosquamosa and Cui et al. (2018) |
| *Amanita clarisquamosa*<br>(*Amanitopsis clarisquamosa* S. Imai 1933)<sup>a</sup> | AF024448 | Weiß et al. (1998)<br>Yang, Z. L. & Tulloss, R. E.<br>http://www.amanitaceae.org/?Amanita%20clarisquamosa and Cui et al. (2018) |

<sup>a</sup> The choice for *Amanita avellaneosquamosa* is particularly difficult. Of the 6 specimens included by Cui et al. (2018) under this name, 5 separate clades were found in the present study. Furthermore, the sequence from specimen HKAS 38300, highlighted in http://www.amanitaceae.org/?Amanita%20avellaneosquamosa (Tulloss and Yang 2024), is quite distant from all other *A. avellaneosquamosa*, as already noted in Cui et al. (2018). The accession AF024441 also considered by Tulloss & Yang (2024) is more consensual, but its origin (Yunnan, China) is rather different from that of the type (Japan; incidentally, the same happens with the ncLSU sequence chosen here to represent *A. clarisquamosa*). Thus, the clade hereby chosen as reference for *A. avellaneosquamosa* is named 'cf. *avellaneosquamosa*1', and more generally we caution that the reference sequences listed in this Table 1 are subject to revision.

### Sources and search strategies

#### Named sequences in Amanitaceae.org (Tulloss and Yang 2024)

The GenBank accession numbers for all sequences in section *Amidella* listed in the details tab of each species were collected and used to retrieve from GenBank the full FastA records using the NCBI Batch Entrez interface (https://www.ncbi.nlm.nih.gov/sites/batchentrez last accessed December 2, 2024).

### 5.8S phylogeny

On February 25^th^ 2023 a total of 7,045 sequences under the genus name *Amanita*, containing at least one of the nuclear ribosomal internal transcribed spacers (ITS), were downloaded from the GenBank database, and processed by ITSx (BengtssonLPalme et al. 2013) in the PlutoF platform (Abarenkov et al. 2010), producing a total of 6,445 5.8S sequences that were aligned in UGene (Okonechnikov et al. 2012) and further processed with MEGA (Tamura et al. 2021), using the Maximum Likelihood tree construction method (see Outline of phylogenetic analysis, below). The resulting dendrogram was then searched for species epithets known to belong to section *Amidella*. Three major clades were identified (Table 2), all containing many accessions without known epithets, and the GenBank accessions contained in them were collected, and used to retrieve from GenBank the full FastA records using the NCBI Batch Entrez interface as mentioned above.

**Table 2.** Major clades derived from the 5.8S phylogeny with known *Amidella* species epithets. Only the epithets of accepted species are in italic, and those in double quotes are probably misapplied.

| Clade | Number of<br>accessions | known species epithets (*Amidella*) | non-*Amidella* nearest species<br>epithets (examples) |
| --- | --- | --- | --- |
| I | 53 | *lanigera*, *brunneomaculata*, *floridella*,<br>*curtipes*, *pseudovalens*, *sagittaria*,<br>“*clarisquamosa*”, *rufobrunnescens* | *goossensfontanae*, *proxima*,<br>*cinereoconia*, *fulvopulverulenta*,<br>*yenii*, *hiltonii* |
| II | 81 | *ponderosa*, *pinophila*, *whetstoneae*,<br>*neocaesariensis*, “*volvata*”,<br>“*avellaneosquamosa*”, “*goossensiae*” | *quenda*, *daucipes*, *walpolei*,<br>*ovoidea* |
| III | 30 | “*avellaneosquamosa*”, *parvicurta*,<br>“*volvata*”, “*peckiana*” | *neo-ovoidea*, *preissii* |

A preliminary phylogenetic analysis of the retrieved ITS sequences was made after this, to constitute an outgroup made only from sequences that branched out closer to the known *Amidella* sequences. Those that ended up in the outgroup were all from clade I (Table 2), all from section *Roanokenses*.

#### Affinity with collected sequences

Two strategies were followed: first, using the voucher identifiers, an in-house wrapper script was developed to parse the output of esearch | efetch | xtract from Entrez Direct (Kans 2013) and used to retrieve the corresponding non-ITS records from GenBank (i.e. for the nuclear ribosomal 28S large subunit gene ncLSU, for the second largest protein subunit of the RNA polymerase II *RPB2*, for the translation elongation factor EF-1 alpha *TEF1*, and for the tubulin beta chain *BTUB*) that matched the specimens already available; second, using the BLAST homology algorithm, either in the PlutoF platform for the ITS sequences (including environmental samples), or in the NCBI interface for ncLSU, *RPB2*, *TEF1* and *BTUB*.

#### Additional sequences, summary and database checks

Several other sequences were spotted as they were referred to in specimen observation online platforms such as MushroomObserver.org, iNaturalist.org or MycoMap.com.

The final search took place on December 2, 2024. A total of 237 ITS, 132 ncLSU, 30 *RPB2*, 24 *TEF1* and 11 *BTUB* nucleotide accessions belonging to section *Amidella* are included in this study (Online Resource 1). Excluded GenBank accessions: MK782535 (it contains errors and was suppressed at the submitter’s request), OQ754998–9 (two partial reads of the same specimen that fit badly in the ncLSU phylogeny), duplicates KT779088, OP681731 and OP681759, as well as those of other genes with very little coverage (the ‘etc’ column in Online Resource 1). Redundant environmental sequences in the UNITE database version 10FU (Abarenkov et al. 2024) were not included either. The ncLSU sequences that appeared to be unique (“singletons”) were entered (using the GenBank accession number) in the search page on ARB-SILVA (Quast et al. 2012) to check for their presence in the ncLSU r138.2 database and retrieve quality parameters dependent on ambiguous, repetitive or chimeric features. Except for accession KP012973, all sequences deposited till August 2018 were thus validated; the exception has a long string of ‘N’ that lies upstream from the ncLSU alignment, and it was decided to include it in the present analyses. Later singleton sequences in our ncLSU dataset were absent from the EUKARYOME ‘blacklist’(Table S1 in Tedersoo et al. 2024). A Species Hypothesis code at 1.5% distance level (Kõljalg et al. 2013) was obtained for most of the ITS sequences.

Geographic references for each sequence were obtained primarily from the GenBank records, complemented by further information that could be retrieved from other sources (scientific publications, the reference website amanitaceae.org (Tulloss and Yang 2024) and online platforms).

### Outline of the phylogenetic analyses

Most ITS sequences were processed in the ITSx programme (Nilsson et al. 2010; BengtssonLPalme et al. 2013) in the PlutoF platform (Abarenkov et al. 2010), marking the ‘Concat’ option to obtain the ITS1:ITS2 concatenate. Those that, for being incomplete, were left unprocessed by ITSx, were aligned to the concatenated sequences, placing a gap in the latter that marked the intercurrent 5.8S gene sequence, thus allowing its safe removal.

The decision to use the ITS1:ITS2 concatenate fell out from the surprising heterogeneity of the 5.8S sequences within *Amidella* (Table 2). This hinted at the possibility that the 5.8S gene might distort the phylogenetic analyses based on the ITS region, and although it was later realised that, as expected from the fact that most of the phylogenetic signal is from the spacer regions, the differences of including the 5.8S gene were minimal (not shown), we had two important reasons to prefer the ITS1:ITS2 concatenate: first, to avoid potential inconsistency in the more critical clades; second, to have a phylogeny based purely on intronic sequences, which are in principle not subject to evolutionary constraints that act on the exonic ribosomal sequences, thus enhancing the contrast with the phylogenetic analyses based on the ncLSU.

All applications used for offline analyses were run in Windows 10. Alignments were made in the UGene platform versions 49–51 (Okonechnikov et al. 2012) using the built-in MUSCLE tool version 5 (Edgar 2022), complemented by visual corrections. The alignments were finally trimmed of terminal tracts with low coverage. They can be retrieved in a figshare repository, https://doi.org/10.6084/m9.figshare.26352388.

The main phylogenetic reconstructions were based on the Bayesian Markov chain Monte Carlo (MCMC) algorithms as implemented in the BEAST package version 1.10.4 (Suchard et al. 2018) supported by the BEAGLE acceleration libraries version 4.0.0 (Suchard and Rambaut 2009; Ayres et al. 2019). Alignments were converted in BEAUTi to XML input files using the default parameters, except that the substitution model was GTR+G+I with 5 gamma categories. Convergence of the MCMC chains in the ITS1:ITS2 reconstruction was confirmed in Tracer version 1.7.2 (Rambaut et al. 2018) with all parameters exhibiting ESS > 200 after a 1,000,000 burn-in. Each BEAST output tree file was then analysed with TreeAnnotator using the default parameters to obtain the Maximum Clade Credibility Tree. A few phylogenetic reconstructions were made using MEGA version 12.0.8 (Kumar et al. 2024). Splitstree CE version 6 (Huson and Bryant 2006) was used for Pairwise Homoplasy Index calculations with the default setting (Bruen et al. 2006).

To evaluate which sequences presumably belong to the same species/Operational Taxonomic Units (OTUs), and to allow the application of p-distance thresholds for tentative species, dendrograms were built with MEGA using the Minimal Evolution method with pairwise deletion, p-distances without corrections and uniform rates among sites (no Gamma correction). In the resulting ‘tree sessions’, using the ‘Collapse Nodes’ menu, option ‘By Sequence Difference’, the branches within a p-distance below a given (adjustable) threshold were condensed. The threshold was estimated for the ncLSU dataset using the barcode gap discovery approach (Puillandre et al. 2012), with the option of including phylogenetic trees in the output. The USEARCH software, version 11, was used as an alternative approach for delimiting centroid-based sequence clusters (Edgar 2010), the centroid-representative sequences aligned in UGene and the alignments used for a phylogenetic reconstruction in MEGA.

Except for the MEGA trees, which were drawn by the integrated tree session module, all trees were rendered in the iTOL website (Letunic and Bork 2024) from input files in text format produced by BEAST and ABGD.

## Results

### Clades at subsection and stirp level

The phylogenetic reconstruction based on the ITS concatenate subset suggests that section *Amidella* is subdivided into five main subclades (Figure 1a) that are named ‘Volvatae’, ‘Sagittariae’, ‘Lepiotidae’, ‘Ponderosae’ and ‘Baakani’. This structure is also obtained for the ncLSU subset, except that Volvatae and Sagittariae are nested in a clade (with 57% posterior probability) that contains all Lepiotidae sequences (Figure 1b). The results also indicate further subdivisions with high support: Volvata, Peckiana and Fulvisquamea (in the Volvatae clade), Lepiotoides, Avellaneosquamosa, Subviscosa and Pseudomutabilis (Lepiotidae clade), Ponderosa, Claristriata and Pinophila (Ponderosae clade) (Table 3). However, two clades containing specimens classified in Lepiotoides and Subviscosa for the ITS concatenate dataset form separate branches in the phylogeny based on the ncLSU dataset.

**Fig. 1.**
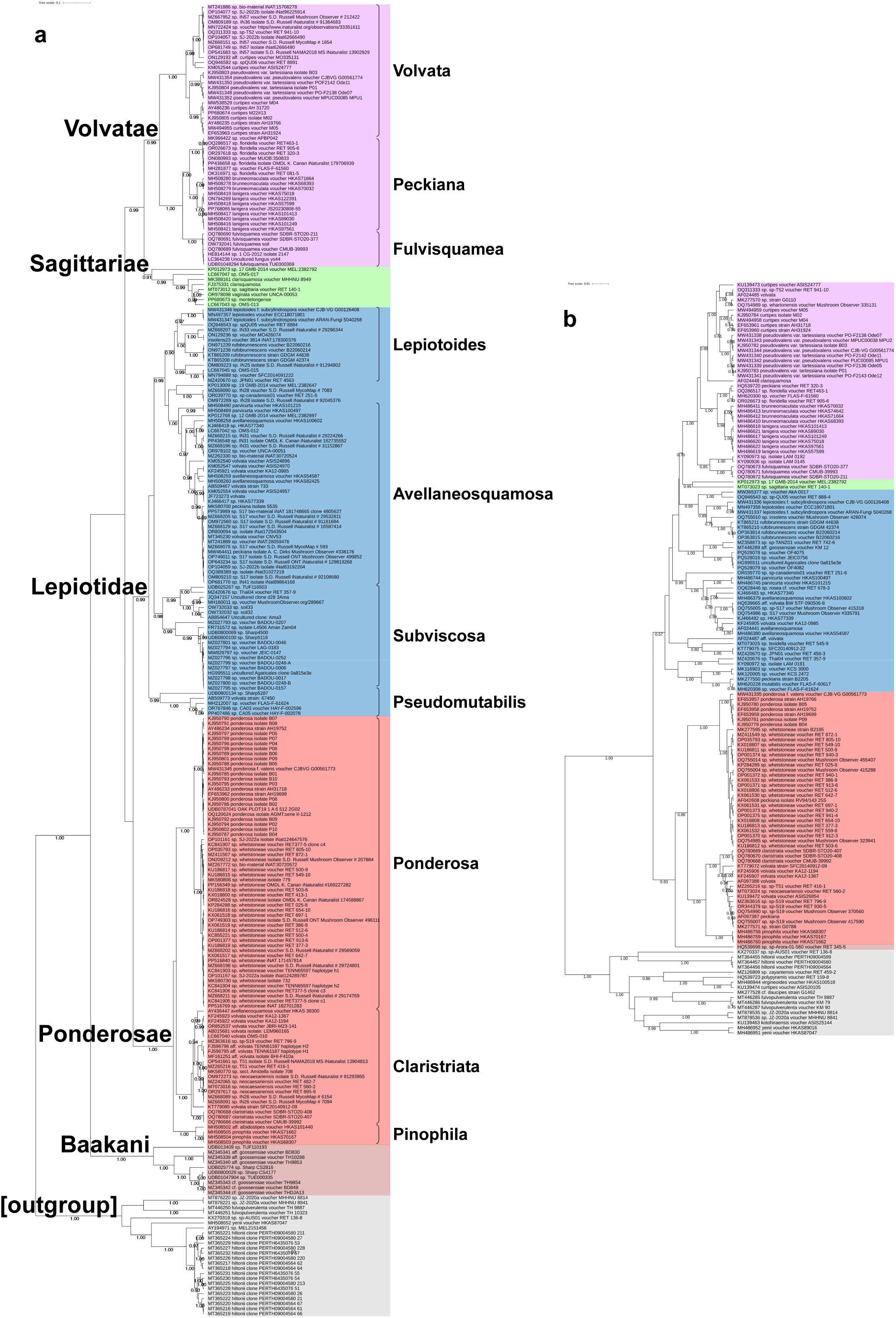
Phylogenetic trees obtained with BEAST (Suchard et al. 2018), with posterior probabilities above 50% shown to the left of each node. The sequence labels have the GenBank accession numbers followed by unique identifiers, and the proposed clades are highlighted with background colours. The scale bars (top left) are for substitutions per site. a) ITS1:ITS2 concatenate dataset, tree log likelihood –14,030.9, with the names for major clades (subsection level, corresponding to a different background colour) and, in smaller type, the minor (stirp level) clades. b) ncLSU dataset, tree log likelihood –7,311.5, background colours matching those of a)

**Table 3.**
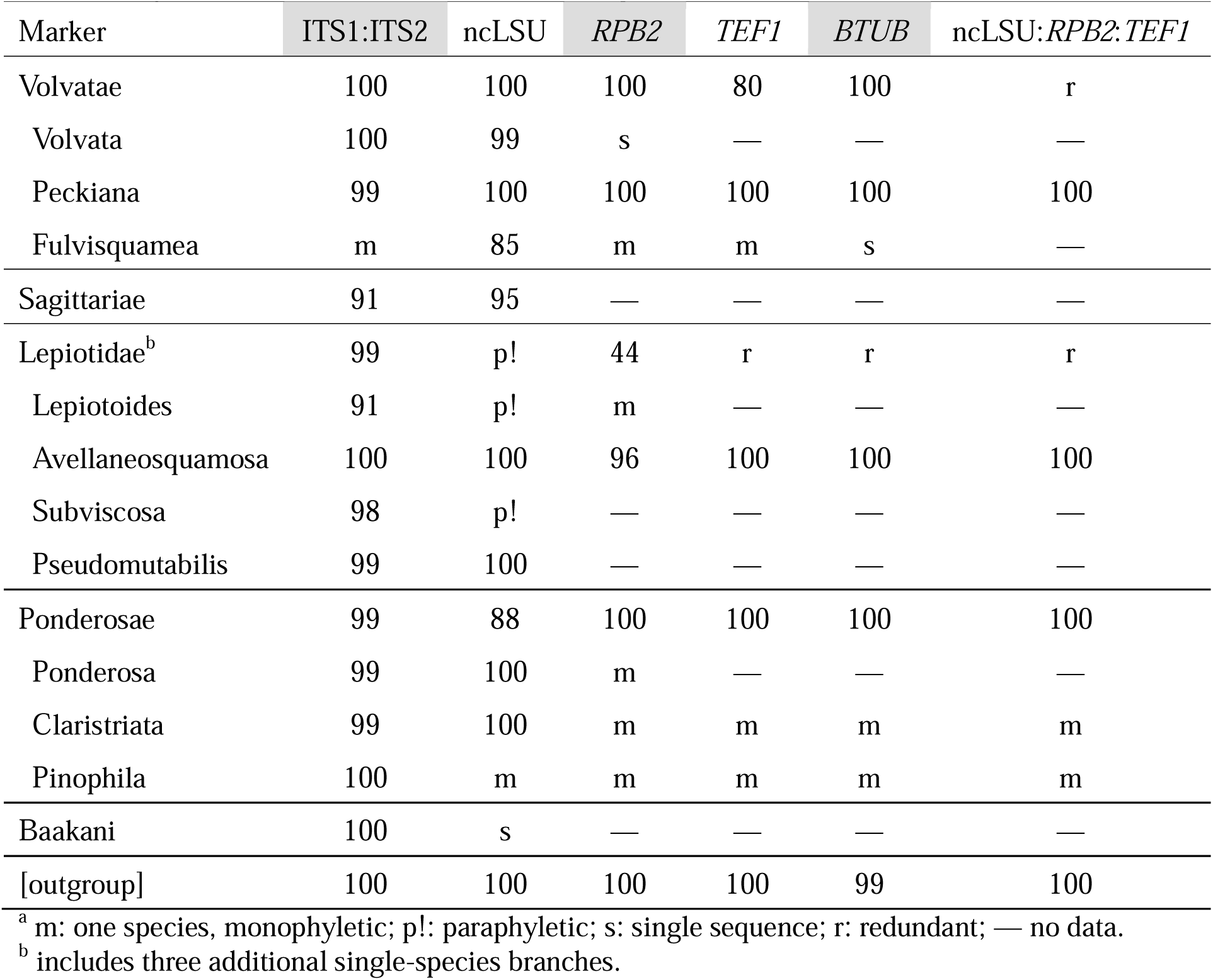
Bayesian posterior probabilities (percentage) for the clades^a^.

Further analyses of other markers (*RPB2*, *TEF1*, *BTUB*) as well as a concatenated ncLSU:*RPB2*:*TEF1* dataset, though limited by the reduced taxa coverage, support this structure (Online Resource 2), the only exception being the low posterior probability for Lepiotidae in the *RPB2* dataset (Table 3). Some of the 100% support values are based on a very limited taxon sampling.

Pairwise Homoplasy Index calculations showed no evidence of recombination, with Φ_w_ > 0.05 in all alignments (Bruen et al. 2006).

### Clades at species level

A phylogenetic reconstruction based on the ITS concatenates was made without distance corrections (i.e., based on the percentage divergences calculated pairwise, and without gamma corrections for between-site heterogeneities) to allow the application of p-distance thresholds for tentative species. By adjusting the minimum distance threshold to separate species at different levels, the 1.25% level was chosen based on the observation that none of the previously recognised species were merged and very few (*Amanita rufobrunnescens*, *A. pseudovalens* and *A. hiltonii*) were split. As discussed in the respective Taxonomy sections below, the *subviscosa* s.l. branches are considered to be three species, while the sp. ‘Naletali’ branch merges two SH codes. At this level, a total of 67 branches were obtained within *Amidella* (Online Resource 3a). For the ncLSU dataset the choice of a 0.8% threshold was determined by the Automatic Barcode Gap Discovery method (Puillandre et al. 2012) and applied as in the ITS concatenate dataset (Online Resource 4). Although in this case a few mergers occur (for example, between *Amanita lanigera* and *A. brunneomaculata*), ad-hoc attempts at lower levels were judged less satisfactory because more taxa were split. The 0.8% threshold value was also applied for the UCLUST clustering algorithm in USEARCH (Edgar 2010) to provide an alternative phylogeny-independent delimitation at species level. Both ABGD and UCLUST provided data for phylogenetic reconstructions that differed only slightly from the one in Online Resource 3b, actually sorting out some of the merged clades in the latter (Online Resource 4).

The ncLSU dataset allowed the identification of the branches that likely correspond to *Amanita volvata*, *A. peckiana*, *A. curtipes*, *A. avellaneosquamosa* and *A. clarisquamosa* (Table 1) and suggested further species not represented in the ITS1:ITS2 dataset. Thus, after studying the possibility of assigning such branches to species already delimited in the latter dataset (see Discussion), a complete list of 81 tentative species was obtained (Table 4). The potential splitting of the two sp. ‘Naletali’ sequences and the potential merging of aff. volvata8 with the sp. S19/aff. volvata9 clade may cause a slight alteration of the total number (see Taxonomy section). Overall, 49 clades at species level (60% of the total of 81) are represented by a unique specimen collection or environmental sequence, and designated “singletons”.

**Table 4.**
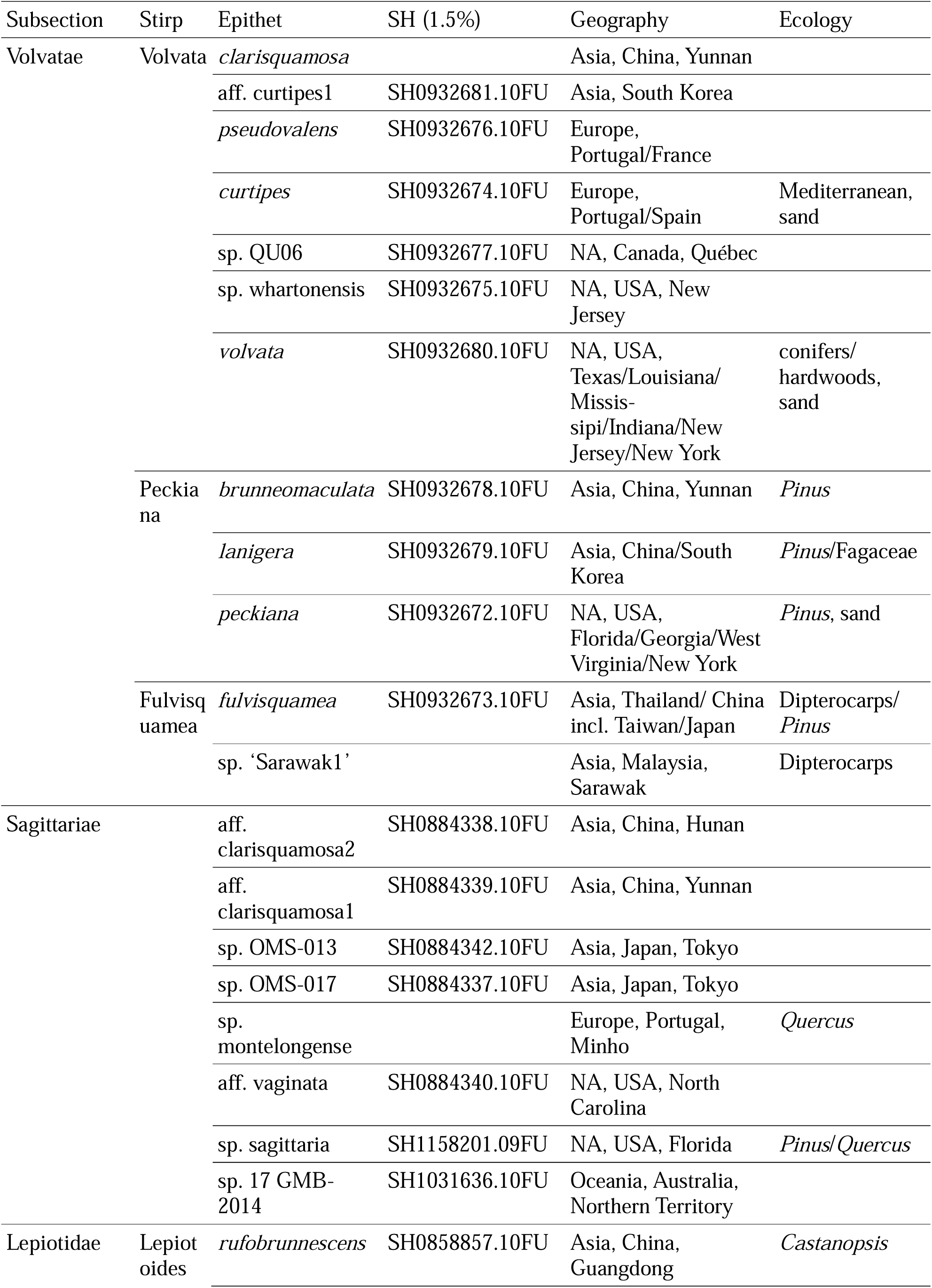
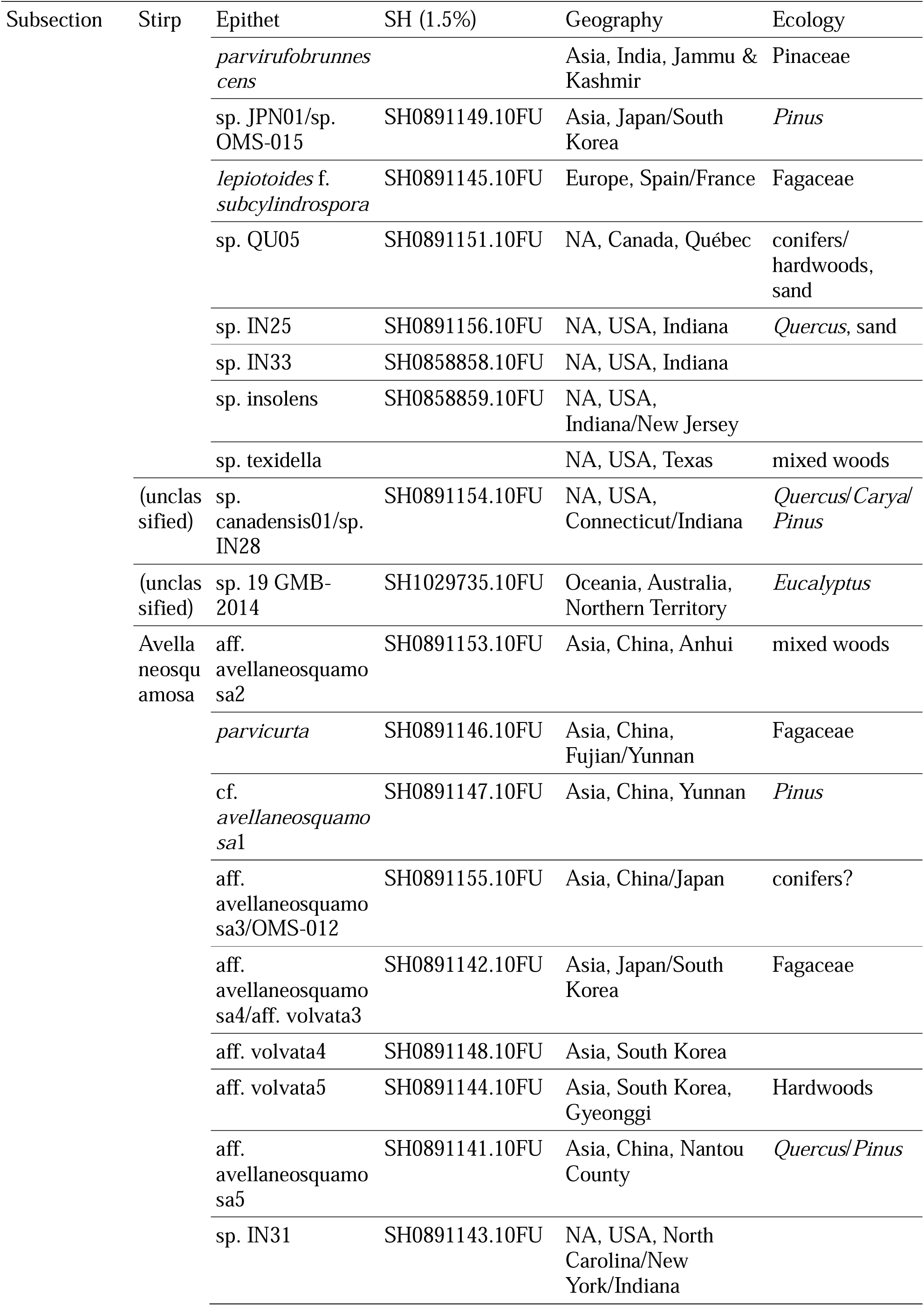
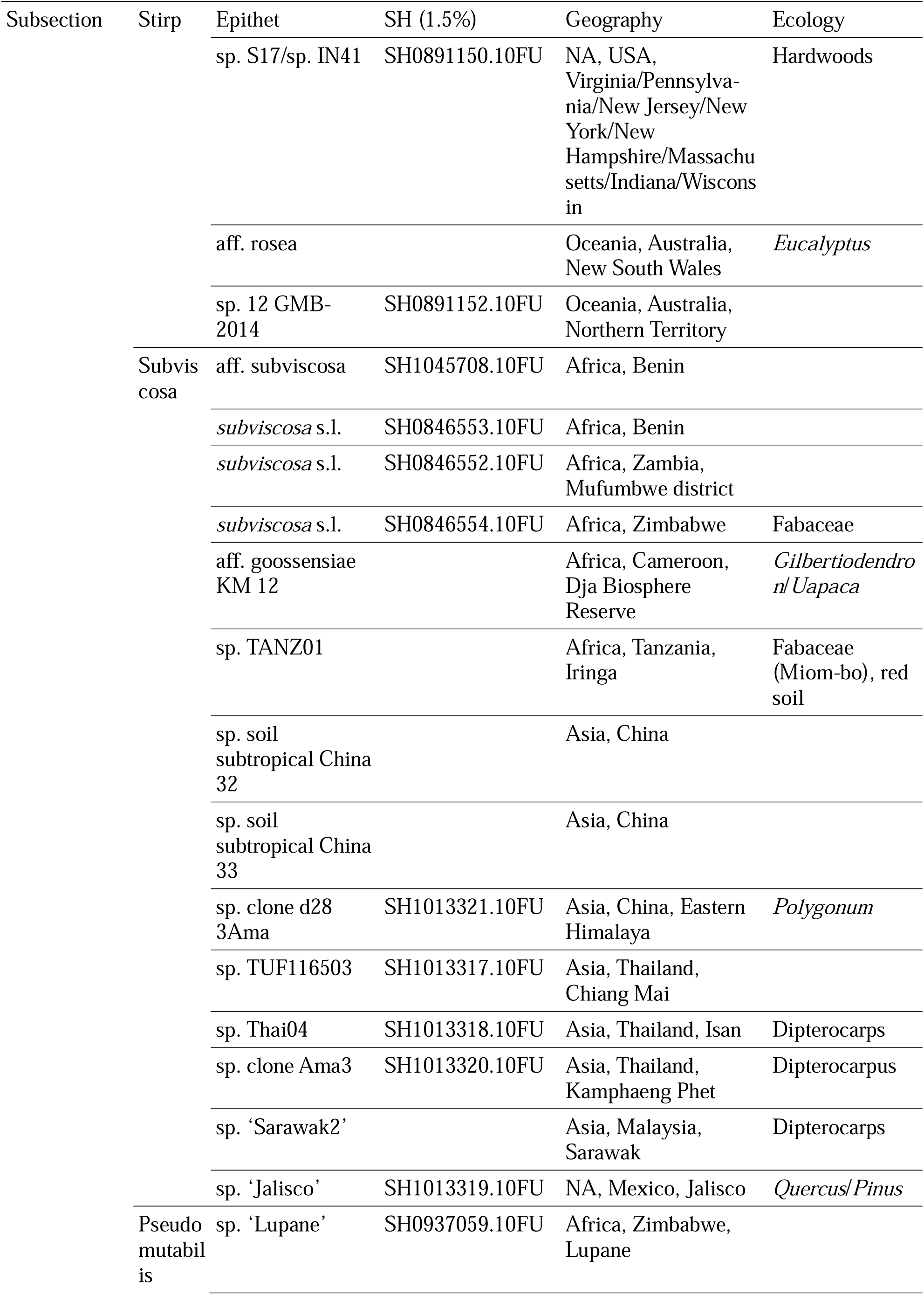
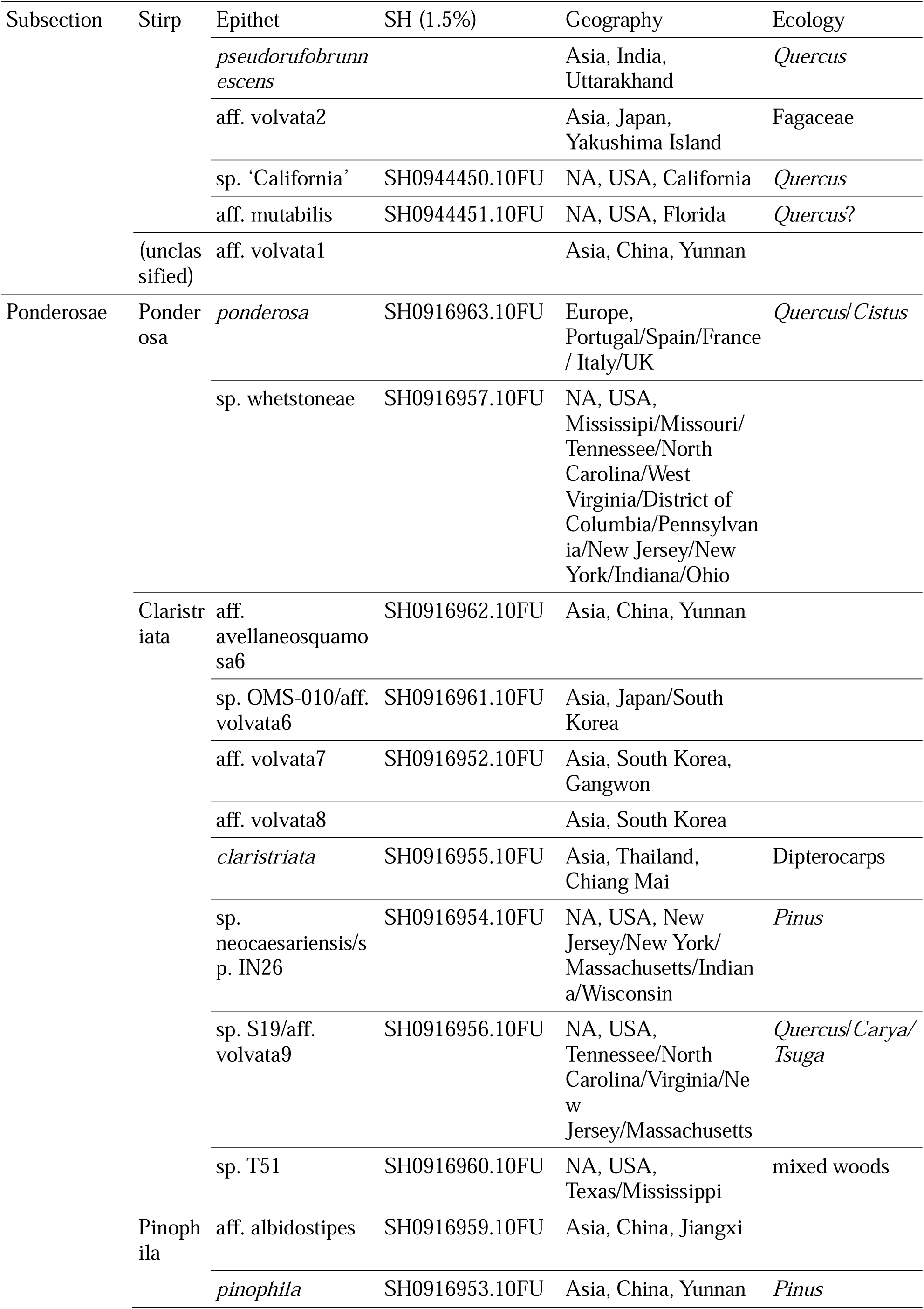
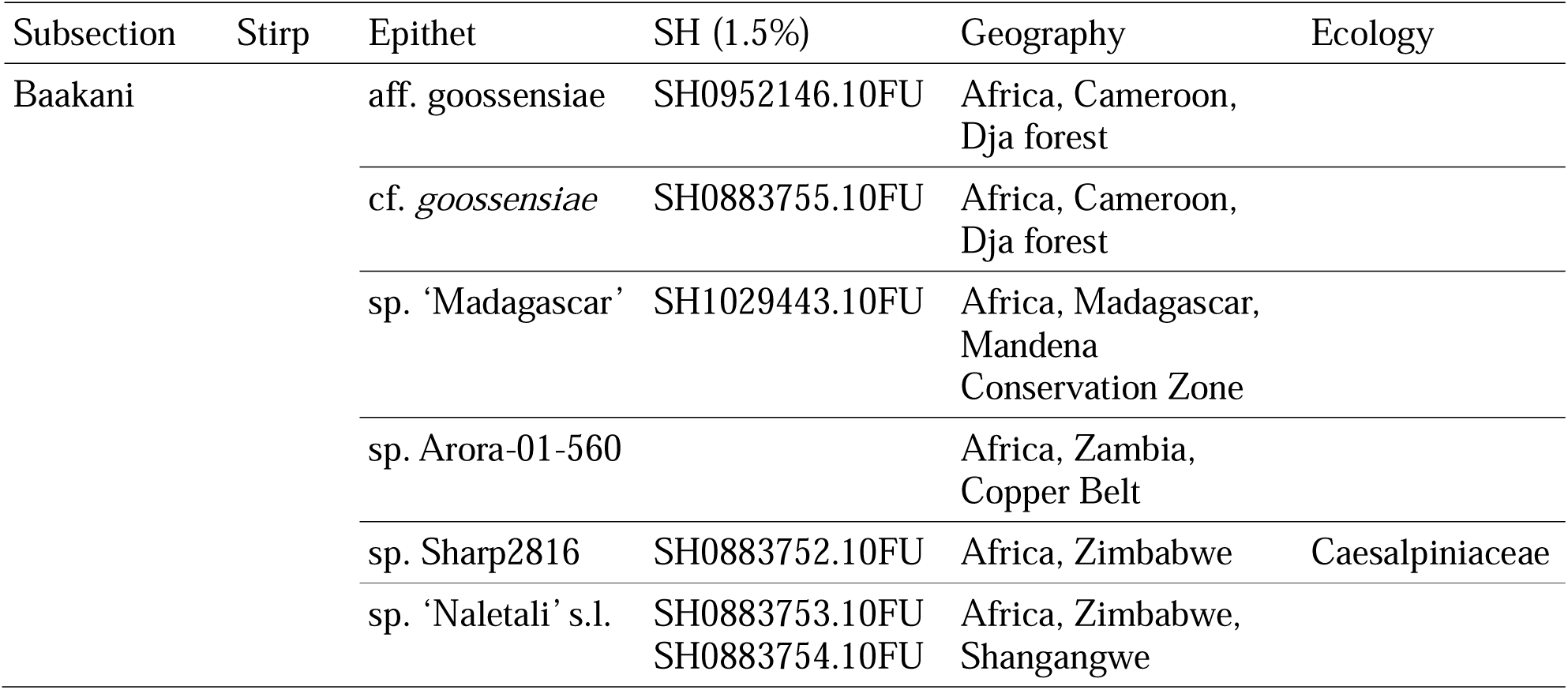
List of the clades at species level, arranged according to the adopted taxonomy (subsection and stirp levels) and alphabetically ordered by geographic location name.

### Biogeography of *Amidella*

Though limited by the geographically uneven representation, the present species outline (Table 4 and Online Resources 5–6) brings out noteworthy aspects. The species are distributed over temperate, subtropical and tropical regions only, comprising 12 of the biogeographic regions defined by Loidi and Vynokurov (2024); the highest diversity is in Asia (40–41 species, half of the total), followed by North America (21 species), Africa (13–14 species), Europe (5 species) and Oceania (4 species). No sequences are known from South America, and 2 species are known (as basidiomes) to be shared between Southern Europe and North Africa (Neville and Poumarat 2004). All Baakani species are from Sub-Saharan Africa and Madagascar.

Furthermore, most species have a limited geographic range. Examples with a relatively wider distribution are *Amanita fulvisquamea*, *A. volvata*, *A. ponderosa* and *A. whetstoneae*, but at least in two of these the ranges might outsize that of the basidiomes: the presence of *A. ponderosa* in oak ectomycorrhizas from South England, for example, is far to the North from the known northernmost occurrence of this species as basidiomes (Neville and Poumarat 2004), and *A. fulvisquamea* basidiomes are known only from Northern Thailand (Liu et al. 2023). Currently, only *A. ponderosa* and *A. pseudovalens* overlap biogeographical regions.

Finally, Tulloss and Yang (2024) list 14 species (some with provisional names) without available sequences (Online Resource 6): seven from Africa (*Amanita floccosolivida* Beeli, *A. fulvosquamulosa* Beeli, *A. goossensiae* Beeli, *A. irreperta* nom. prov. E.-J. Gilbert, *A. olivacea* Beeli, *A. subviscosa* Beeli, *A. subviscosa* sensu Pegler & Shah-Sm.), one from Asia (*A. duplex* Corner & Bas), four from North America (*A. dolichopus* nom. prov. Tulloss, *A. fallax* nom. prov. Tulloss & G. Wright, *A. occidentalis* O. K. Mill. & D. J. Lodge, *A. pseudovolvata* nom. prov. Tulloss) and three from Oceania (*A. curta* (Cooke & Massee) E.-J. Gilbert, *A. grisea* Massee & Rodway, *A. pallidogrisea* A. E. Wood). Given that *A. floccosolivida* and *A. pallidogrisea* may not belong to section *Amidella* (Tulloss and Yang 2024), it would be less likely to have them captured in our search; apart from that, a relation to specimens or samples for which any of the classified sequences are known seems to be possible only for *A. subviscosa* sensu Pegler & Shah-Sm. (noting in particular the Zambian *subviscosa* s.l. species clade), *A. irreperta* (to the sp. ‘Madagascar’ clade) and *A. fallax* (most likely the sp. ‘California’ clade).

## Taxonomy

We first propose an update of the description of section *Amidella* and later outline the species composition of the clades within *Amidella* (Table 4).

### Updated description for section *Amidella*

In their taxonomic scheme for European species of *Amanita*, Neville and Poumarat (2004) adopted subsection *Ovoideinae* Singer, corresponding to the original genus *Amidella* E.-J. Gilbert, and within it they made a split between series *Ovoidea* and series *Amidella*, proposing for the first time to our knowledge a separation between the current concept of *Amidella* and species that have been replaced in section *Roanokenses* (Wolfe et al. 2012; Cui et al. 2018; Riccioni et al. 2019). The keys to their *Amidella* series result in the following diagnosis (our translation from the original in French):

Margin of the pileus smooth and sometimes appendiculate, saccate volva under the stipe, annulus rudimentary or absent, quick discolouration of the context to reddish tones either by surface rubbing or exposure to air, later to brownish tones, amyloid spores (Neville and Poumarat 2004).

According to Neville and Poumarat (2004), the separation of the two series is for “reasons of convenience” based on the discolourations. However, for non-European species the above diagnosis and the discolouration criterium require further clarification: besides the fact that in many specimens the pileus margin is striated and not smooth (Bas 1969), the requirement for discolouration is left to some questioning, with an “unchanging trama” commonly reported, while several species in the current concept of *Roanokenses* have discolourations considered typical of *Amidella* — for example *Amanita daucipes* (Mont.) Lloyd, *A. fulvopulverulenta* Beeli, *A. luteofolia* Yang-Yang Cui, Qing Cai & Zhu L. Yang or *A. mutabilis* Beardslee (Tulloss 1984; Cui et al. 2018; Tulloss and Yang 2024). It is only in combination with other characters, namely the saccate volva and lack of a persistent membranous ring, that the discolourations can help define *Amidella* (Online Resource 7).

Currently, at section level, *Amidella* is the taxon introduced by Konrad and Maublanc (1948), who sweepingly transferred it from the one defined by Gilbert at subgenus level (Bas 1969). Although the taxon is not listed in the nomenclatural databases Indexfungorum (Royal Botanic Gardens Kew Mycology Section et al. 2024) and Mycobank (Robert et al. 2005), it is considered valid by different authorities (Neville and Poumarat 2004; Cui et al. 2018; Tulloss and Yang 2024). However, the concept of section *Amidella* needs revision because, contrary to Gilbert’s nuanced original description of the protolog for genus *Amidella* (Neville and Poumarat 2004), it excludes striation of the pileus margin, which is sometimes mentioned for the section (Bas 1969; Yang et al. 2001; Sanmee et al. 2008) and specifies a bulbous stipe base (Konrad and Maublanc 1948). Furthermore, we find that several European species show a faint greenish-grey reaction with iron sulphate on the exposed context, as suggested first by the analysis of *Amanita ponderosa* (Tulloss and Yang 2024), and in contradiction to a rule put forward for the whole genus (Neville and Poumarat 2004). We believe that this reaction should be systematically tested for the whole of *Amanita* in order to determine its phylogenetic extent. Therefore, we propose the following update:

### Section *Amidella* Konrad & Maublanc mut. char. (emend. P. Oliveira & R. Arraiano-Castilho)

#### Type species: *Amanita volvata* (Peck) Lloyd

Agaricoid basidiomata, all parts initially whitish and becoming ochre to tan, sometimes more intensely in the lamellae, with aging or drying. Usually, the surface of fresh specimens quickly reddens where rubbed or bruised, in some cases with a brownish or greyish discolouration. Universal veil persisting as a saccate membranous volva around the stipe base and, sometimes, as membranous patches on the pileus, with remnants of the inner friable layer on the lower part of the stipe and/or on the pileus standing out as earlier and more intensely coloured markings, forming zebroid patterns on the stipe and/or squamules or patches (lepiotoid appearance) on the pileus. Remains of the partial veil adherent to the upper part of the stipe, sometimes contrasting with the more intensely discoloured friable remains of the universal veil on the lower part, forming in some immature specimens an evanescent creamy annulus and/or tiny appendiculate rags on the pileus margin. The pileus margin can be striated (*Vaginatae* appearance). Lamellulae truncate to nearly truncate, lamellar edge normally concolorous. Stipe base usually not bulbous in cross section (discounting the added thickening by the volva). Context (fresh) frequently showing a quick reddening on exposure, and/or (fresh as well as dried) a faint grey reaction with iron sulphate solution. Spores amyloid (exceptionally inamyloid), usually elongate to cylindric (typically average Q > 1.6), septa without clamps in all parts.

### Clades at subsection level

#### Volvatae

The proposed name assumes that *Amanita volvata*, the type species for section *Amidella*, is represented by accession AF024485 (Table 1). At least one close ncLSU sequence (GenBank accession OQ311333) is from a specimen also represented in the ITS concatenate dataset, thus enabling the merging of various ITS sequences within this species and forming the reference species clade for section *Amidella*.

Neville and Poumarat (2004) have used *Volvatae* at section level, following previous authors to encompass (for Europe) current sections *Amidella*, *Phalloideae*, *Validae* and part of *Roanokenses*, thus it is an altogether different taxon.

Three resolved and generally well-supported subclades at stirp level are considered:

1. Volvata Comprises the type species *Amanita volvata* (Peck) Lloyd, *A. clarisquamosa* (S. Imai) S. Imai, *A. curtipes* E.-J. Gilbert (all based on the assignments in Table 1), *A. pseudovalens* (Neville & Poumarat) Arraiano-Castilho et al., and 3 unnamed or provisional species (Table 4). According to our analyses, a variety of provisional names from the Eastern United States (sp. T52 Tulloss & S. D. Russell, IN57, IN36) are all *A. volvata*.
2. Peckiana Comprises *Amanita peckiana* Kauffman in Peck (comprising sequences from specimens clustering around accession HQ539720, Table 1), *A. brunneomaculata* Yang-Yang Cui, Qing Cai & Zhu L. Yang, and *A. lanigera* Yang-Yang Cui, Qing Cai & Zhu L. Yang (Table 4). According to our analyses, *A. floridella* Tulloss & S. D. Russell nom. prov. is *A. peckiana*.
3. Fulvisquamea Comprises *Amanita fulvisquamea* Yuan S. Liu & S. Lumyong and *A*. sp. ‘Sarawak1’, the latter represented by ncLSU sequences clustering with those of *A. fulvisquamea* with moderate support (Table 3).

#### Sagittariae

This clade is composed of 8 singletons, and named after *Amanita sagittaria* Tulloss & K. W. Hughes nom. prov.. Two sequences are named *A. clarisquamosa*, but the specimen assumed to represent this species (Table 1) has an ncLSU sequence in the Volvatae clade, unrelated to Sagittariae specimens (Figure 1b); therefore, under the assumption of Table 1 the *clarisquamosa* epithet may not be applicable in clade Sagittariae.

#### Lepiotidae

The proposed name is based on the oldest accepted epithet present, *Amanita lepiotoides*. This group of species is by far the most diverse (about half of section *Amidella*). Notwithstanding the strong support in the ITS concatenate phylogeny for this clade and its main subclades (Figure 1a), the ncLSU phylogeny shows the corresponding group in a clade with low support that includes Volvatae and Sagittariae (Figure 1b, Table 3), and with an even lower support in the *RPB2* phylogeny but separate from Volvatae (Table 3, Online Resource 2a). Therefore, the supposed natural groups in the ITS phylogeny should be viewed with caution.

Moreover, there is a potential difficulty with the names to be used. The available sequences named *Amanita lepiotoides* are of the forma *subcylindrospora* (Neville and Poumarat 2004; Arraiano-Castilho et al. 2022) and one cannot be certain whether the autonym form would be phylogenetically close or not. Furthermore, as pointed out in the footnote to Table 1, the original *A. avellaneosquamosa* could be different from our assumption, noted here as ‘cf. avellaneosquamosa1’ (Table 4, Online Resources 1 and 3). Hence, if any of these two taxa turn out to belong outside the clades under consideration, the respective naming should be revised.

Based on the ITS concatenate phylogeny, four subclades are considered at stirp level:

1. Lepiotoides Comprises *Amanita lepiotoides* f. *subcylindrospora* Neville & Poumarat, *A. rufobrunnescens* W. Q. Deng, T. H. Li & Hosen, *A. parvirufobrunnescens* A. Kumar, Y. P. Sharma & Mehmood, *A. insolens* I. Safonov & L. V. Kudzma nom. prov., and up to 4 other species (Table 4). One of the latter might be *A. texidella* Tulloss, Pastorino & K. W. Hughes nom. prov., since its ncLSU sequence is close to *A.* sp. JPN01/sp. OMS-015 (80% posterior probability), but the branch containing this species pair is separate from the remainder of the Lepiotoides sequences.
2. Avellaneosquamosa Comprises 12 clades, including *Amanita parvicurta* Yang-Yang Cui, Qing Cai & Zhu L. Yang and the specimen presumed to represent the original *A. avellaneosquamosa* (S. Imai) S. Imai in Ito, included here in clade *A.* cf*. avellaneosquamosa1* (see note on Table 1), as well as 10 unnamed species. The epithet *rosea*, given with a cf. to one ncLSU sequence (accession OQ628446), is of a species belonging in section *Roanokenses*, and for this reason *A.* aff. *rosea* is used to name the corresponding clade.
3. Subviscosa This branch bears the name of *Amanita subviscosa* Beeli but this species might not be represented in the current dataset: a group of three closely related species clades, each corresponding to geographically segregate SH codes, was named *A. subviscosa* s.l. instead, for it is uncertain whether any of these correspond to the described *A. subviscosa* (Table 4, see also Discussion). The Subviscosa group comprises up to 14 species from warm climates, including almost all from Africa that are not Baakani (Table 4). Although the ITS phylogeny places *A.* sp. Thai04 in this group, this species forms a separate branch in the ncLSU phylogeny, together with the additional *A.* sp. ‘Sarawak2’, thus the inclusion of both in Subviscosa is rather tentative. The ncLSU sequence named *A.* aff. goossensiae KM 12 has a position in this subclade (close to *A. subviscosa* s.l.) that appears not to be the one expected for the clade Baakani (see below), where other specimens from the same Cameroon location are also named in approximation of the same epithet.
4. Pseudomutabilis This clade is named in connection with one of the ncLSU sequences (accession MH620228) identified as *Amanita mutabilis*, which is expected to be wrong since this is a species with a membranous ring belonging in section *Roanokenses* (Tulloss & Yang 2024). The species clade for that accession is named *A.* aff. *mutabilis*. Further species clades include *A. pseudorufobrunnescens* K.C. Semwal, K. Das, R.P. Bhatt, Mehmood & V.K. Bhatt, and three others, one of them possibly *A. fallax* nom. prov. as noted in the Biogeography section above.
5. Unclassified There are three Lepiotidae species that form single branches and thus are kept unclassified: *Amanita* sp. 19 GMB-2014 and *A.* sp. canadensis01/sp. IN28 are placed next to Avellaneosquamosa (Figure 1a-b); and *A.* aff. volvata1, represented by a single ncLSU sequence.

#### Ponderosae

The proposed name is based on the oldest accepted species in this clade, *Amanita ponderosa*. Three well-resolved and well-supported subclades at stirp level are considered:

1. Ponderosa Comprises *Amanita ponderosa* Malençon & R. Heim and *A. whetstoneae* Tulloss, Goldman and Kudzma nom. prov..
2. Claristriata Comprises *Amanita claristriata* Yuan S. Liu & S. Lumyong, *A. neocaesariensis* Tulloss, I. Safonov & K. W. Hughes nom. prov. (including specimens labelled IN26), and around 6 other unnamed species. The uncertainty in the number of species is due to the limited species-level correspondence between some ITS clades (sp. OMS-010/aff. volvata6, aff. volvata7, aff. volvata8, sp. S19/aff. volvata9) and their presumed ncLSU counterparts (see Discussion).
3. Pinophila Comprises *Amanita pinophila* Yang-Yang Cui, Qing Cai & Zhu L. Yang and an unidentified specimen from China labelled *A.* aff. *albidostipes*, an epithet that applies to a species in section *Vaginatae* (Cui et al. 2018).

#### Baakani

This clade is named with reference to the Ba’Aka nation in Cameroon, who live in the provenance of six undescribed specimens named for the apparent affinity to *Amanita gossensiae* Beeli, a species classified in section *Amidella* (Tulloss and Yang 2024). These specimens belong in 2 species, and another 2–3 species from Zimbabwe and 1 from Madagascar also belong in this clade. Furthermore, an unnamed species from Zambia (*A*. sp. Arora-01-560) represented by a single ncLSU sequence has a position in the phylogeny congruent with belonging in Baakani. We note that the position of *A. goossensiae* still needs to be confirmed, and a separate collection named *A.* aff. *goossensiae* KM 12, from the same Cameroon provenance, might be an alternative candidate, but its ncLSU sequence is placed within Lepiotidae (see Subviscosa, above; Figure 1b).

#### Hesleri

Sequences belonging to either *Amanita hesleri* Bas or *A. zangii* Zhu L. Yang, T.H. Li & X.L. Wu, which belong to section *Phalloideae* (Cai et al. 2014; Cui et al. 2018), frequently appeared in the BLAST results, and early phylogenetic reconstructions of *Amidella* suggested them to form a clade intermediate between the outgroup and *Amidella*, even forming a clade with the Baakani sequences in one case (not shown), thus raising questions about the actual limits between sections. Their morphologies are atypical for *Phalloideae* (Cui et al. 2018) and certainly do not conform to the established concept of section *Amidella*, namely for the lack of saccate volva and lack of discolourations of the basidiome, as well as for the microscopy of the volva remnants on the pileus (Yang et al. 2001; Tulloss and Yang 2024), so their placement had to be examined, both for the ITS1:ITS2 concatenate and the ncLSU. The results confirmed their position within *Phalloideae*, neatly separate from *Amidella* (Online Resource 8). The short branch separating Hesleri from the *Phalloideae* node in most phylogenies may explain why these taxa were retrieved in our BLAST-based searches.

## Discussion

The present work collects for the first time a comprehensive coverage of the sequence data available in section *Amidella*, establishing its limits with regard to neighbouring sections, delimiting clades at species level and thereby tripling the total count, uncovering their arrangement into subsectional clades and opening a new discussion on their biogeography.

### Evidence for subsectional clades and undescribed diversity

The present work presents for the first time, as far as the authors are aware, a nearly exhaustive sampling of the DNA sequence diversity available for section *Amidella*. Our analyses provide a boundary separating the section from neighbouring sections *Roanokenses*, *Arenariae* and *Phalloideae*, to include unsuspected phylogenetic branches even in the face of misleading epithets such as vaginata. Furthermore, a previously unknown structure is revealed, with five clades at subsection level, with further subdivision at stirp level in three of them. This structure adds at least one level of precision in the currently problematic task of species determination, and can be very useful in collecting sequences for phylogenetic analyses more effectively, both by overriding the need for very large datasets in studies of closely related groups (Dissanayake et al. 2024) and by helping form a balanced representation of the groups to be studied. Although well-supported in general (the main exception is a partial disruption of Lepiotidae in the ncLSU phylogeny, Figure 1b, Table 3), the topology between the five clades is rather constant, with Baakani as the early-branching clade, while Sagittariae and Volvatae appear as the most derived. It remains to be seen whether it will hold up with the publication of new taxa, for example the five vouchers from China (Cai et al. 2024) placed within *Amidella* thanks to the *TEF1* sequences released so far (HKAS57709/69526/84822/92026/129313, Online Resource 9).

The complexity within *Amidella* is also expressed by a total of about 81 clades at species level (Table 4, Online Resource 3), of which only 16–17 have accepted names (Table 4). Adding 9–12 described species without sequences (Online Resource 6), the current total for section *Amidella* reaches circa 90 species, of which only one third have accepted names. The delimitation of clades at species level was done with a branch clustering option in MEGA, using consensus cutoffs for the pairwise uncorrected p-distances that, in this group, were set at 1.25% for the ITS concatenate and 0.8% for the ncLSU. This approach is tree-based and is not limited by the high prevalence of singletons, which is an obstacle for statistical approaches that rely on intraspecies diversity such as sGMYC or mPTP (Fujisawa and Barraclough 2013; Leavitt et al. 2015; Sánchez-Ramírez et al. 2017; Maharachchikumbura et al. 2021). However, the cutoff used for a given phylogeny is likely heterogeneous in different branches, as suggested by the slight differences between different approaches to the ncLSU dataset (Online Resources 3b and 4). Notwithstanding the suggestion that only the ITS2 region has an effective barcoding gap for *Amanita* (Badotti et al. 2017), the high prevalence of species not represented by ITS sequences argues for the use of all information available. At any rate, the evidence of a high number of novel species, mainly in clades Baakani, Sagittariae, Lepiotidae (subsection level) and Claristriata (stirp level), is compelling.

It is worth mentioning that, because of nrDNA cistron heterogeneity, a single sequence might not be representative of the collection itself — as so well exemplified in Online Resource 3a with the *Amanita hiltonii* ITS sequences in the outgroup (Davison and Giustiniano 2020). Genomic analyses have demonstrated that the same can happen in the ncLSU (Paloi et al. 2022; Cedeño-Sanchez et al. 2024). Nevertheless, a recent survey has obtained a >99% pairwise identity between ITS cistrons in *Amanita* (Bradshaw et al. 2023), therefore we feel that nrDNA cistron heterogeneity would affect, if anything, the separations at interspecies level rather than higher-level relationships.

Unfortunately, DNA sequences are the only support for these subclades. Even with the very detailed descriptions available (Deng et al. 2016; Cui et al. 2018; Semwal et al. 2020; Liu et al. 2023; Tulloss and Yang 2024), attempts to derive morphology-based infrasectional simplesiomorphies were unsuccessful (not shown; Online Resource 7 shows examples of how some variations on the overall morphological unity of the section, as outlined in our proposed description, are shared by more than one infrasectional clade). This, and the possibility of minor changes as more species are discovered, deters their formal proposal as infrasectional taxa. On the other hand, the available alignments suggest efficient means to design taxon-specific PCR primers to retrieve enough sequence information from type specimens and thus resolve the hitherto unclear identification of taxa such as *Amanita avellaneosquamosa* (Table 1; Online Resource 10).

### Biogeographical implications

Strikingly, practically all species have a relatively narrow geographical range (Table 4, Online Resource 5), as reported already for other taxa (Looney et al. 2016, 2020; Peay and Matheny 2016; Bazzicalupo et al. 2019; Noffsinger et al. 2024). This could help avoid using the same name for geographically distinct collection origins — as the clade naming used here at species level demonstrates, the *volvata*, *peckiana*, *curtipes* and *avellaneosquamosa* epithets have been used very confusingly (Figure 1, Table 4, Online Resource 1). Four contrasting examples serve to highlight the application of this geographic premise, as well as difficulties that may arise:

i. The clade *Amanita subviscosa* s.l. spans an unusually vast distribution in Africa, from Benin to Zambia and Zimbabwe (Table 4). We named it so for two reasons: first, the SH codes at 1.5% are unique for each country, thus suggesting that three species of narrower range are to be considered; second, *A. subviscosa* has been described from a fourth country (Online Resource 6) and is regarded as different from an unsequenced collection from Zambia (Tulloss and Yang 2024), therefore one cannot be sure which of the sequenced specimens, if any, represents *A. subviscosa*.
ii. There are three North American ITS clades in Claristriata (Ponderosae) with contiguous geographical distributions: *Amanita* sp. S19/aff. volvata9 ranging from Massachusetts to Tennessee, *A.* sp. neocaesariensis/sp. IN26 from New York to Wisconsin, and *A.* sp. T51 in Mississippi and Eastern Texas. The latter two collapse into a single lineage at 1.4% divergence, raising the possibility that they could be a single species, but their disjunct distributions suggest vicariant species. The three clades merge at 1.55% divergence, strengthening the hypothesis of a recent speciation. They might be useful for a case study of a speciation process in ectomycorrhizal fungi.
iii. Still with the *Amanita* sp. S19/aff. volvata9 group, there is an ncLSU sequence from a South Korean collection that is very close phylogenetically. An ITS sequence for this collection would help understand its relationship with the North American group, but none is available. Tentatively, we consider the South Korean collection autochthonous and assign a separate epithet (volvata8), but it could turn out to be the same species as the North American collections, and one possible explanation for such a wide distribution could be the accidental introduction with an ectomycorrhizal host originating in the United States (Kim and Zsuffa 1994; Vellinga et al. 2009).
iv. The provenances of Eastern Asia (specifically, China — Mainland and Taiwan — Japan and South Korea) have only a minority of shared species (Online Resource 11), most of them between Japan and South Korea, while Mainland China shares only 2 out of 16 with Japan (one being *Amanita* aff. avellaneosquamosa3/OMS-012, but the Chinese occurrence is in a botanical garden in Jiangxi; the other is *A. fulvisquamea*, a rather wide-ranging species when soil and root tip samples are considered, however it occurs as fruitbodies only in Thailand until now). One should expect these separations to blur as more findings are reported, but currently, for any species in this group, it is worth considering with caution a shared geographical distribution between Mainland China and Japan, as we note on Table 1.

### Uniqueness of species defined with different datasets

A considerable part of the species deduced from clades in the present work (twelve, to be precise) are represented by ncLSU sequences without an ITS counterpart. This raises two questions: i) is any of these species the same as one of those defined by ITS sequences without ncLSU counterpart, such that both should be merged, and ii) is the species threshold used for ncLSU (0.8% p-distance, Online Resource 3b) anywhere less applicable?

To answer the first question, the cases to be considered for merging narrow down to a few possible pairs classified in the same subclade and possibly not too far geographically: in Claristriata, there are *Amanita* aff. avellaneosquamosa6 (ITS; China, Yunnan) and *A.* aff. volvata8 (LSU; South Korea), but the latter is phylogenetically very close to *A.* sp. S19/aff. volvata9 (Eastern United States) and the former is not; in Avellaneosquamosa, *Amanita* aff. rosea (LSU) is Australian like *Amanita* sp. 19 GMB-2014 (ITS) and both are phylogenetically close to *Amanita parvicurta*, but the first is from New South Wales and the second from Northern Territory; in Lepiotidae, *Amanita* aff. volvata1 (LSU; China, Yunnan) remains unclassified and the species that are nearest geographically (in Thailand and South China) are in unrelated Subviscosa branches; in Baakani, the *Amanita* sp. Arora-01-560 specimen (LSU; Zambia) could be of the same species as one of those in the clade represented by ITS sequences, especially those in Zimbabwe, at least 1000 Km away nevertheless. Hence, each of these four pairs seems unlikely to be merged. Finally, the *Amanita* sp. S17/sp. IN41 clade has a single publicly available ncLSU accession (HQ539665) for the reference specimen RET 444-9 (Tulloss and Yang 2024), but there are several others in the ITS concatenate dataset identified as *Amanita* sp. S17, in a cluster that includes specimens designated *A.* sp. IN41, *A. “*volvata” or *A.* “peckiana”. This extension of *Amanita* sp. S17 seems reasonable since all specimens are from Eastern United States, with similar ecology and phenology and, like accession HQ539665, they cluster with the same group of clades at species level (*Amanita* aff. volvata5, *A.* aff. avellaneosquamosa4/aff. volvata3, *A.* cf. avellaneosquamosa1).

The second question highlights a contrast between the criterium used here for the ITS (1.25%, Online Resource 3a) and the one used for the ncLSU phylogeny (0.8%, Online Resource 3b). While the former was chosen by trial and error, guided by branch collapsing within accepted species, for the latter there was no consensus value that could mimic the results with the species also present in the ITS concatenate dataset. Thus, the ABGD method (automatic barcoding gap discovery) was used to obtain the 0.8% estimate for the p-distance to be used for ncLSU, to settle on a common threshold (Online Resource 3b) and choose the similarity level of 99.2% for UCLUST, used as a phylogeny-independent approach to delimiting sequence clusters. From both approaches it was possible to obtain alternative ncLSU groupings (Online Resource 4) not unlike the one in Online Resource 3b, separating adequately some of the merged clades in the latter, for example the one containing *Amanita lanigera* and *A. brunneomaculata*. We take these results as independent support for the proposed species list (Table 4), but it remains admissible that part of those twelve ncLSU-only clades might be later considered for merging with others, especially as the respective ITS sequences become known.

## Conclusion and Outlook

The present study exposes the unsuspected dimension of some knowledge gaps regarding section *Amidella*: a large majority of undescribed species, a sizeable number of described species without molecular information, the need for molecular studies on types to clarify the identity of species such as *Amanita avellaneosquamosa* and *A. clarisquamosa*, the narrow geographical range as a rule, and an essential need for novel discriminant characters, the utility of which is framed from now on by the infrasectional taxonomic structure here revealed.

One final point. In naming the clades, we had no choice but to use three epithets (*fulvisquamea*, *claristriata*, *pinophila*) that have been accepted only recently (Cui et al. 2018; Liu et al. 2023), *sagittaria* that is still provisional (Tulloss and Yang 2024), and devise a novel name for the Baakani clade. Section *Amidella* is still a largely untrodden field of research.

## Supporting information

Supplemental figures, tables and protocol

## Acknowledgement

Three anonymous reviewers contributed valuable comments on the manuscript.

## Statements and Declarations

### Ethical Approval

Research involving Human Participants and/or Animals: not applicable.

### Consent to participate

Not applicable

### Consent to publish

All authors agreed with the content and all gave explicit consent to submit and they obtained consent from the responsible authorities at the organization where the work has been carried out, before the work is submitted.

### Data Availability Statement

The datasets generated and analysed during the current study are available in the figshare repository through the hyperlink https://doi.org/10.6084/m9.figshare.26352388.

### Authors Contributions

Both authors made substantial contributions to the conception of the work, and to the acquisition, analysis, and interpretation of data; both authors drafted the work and revised it critically for important intellectual content; both authors approved the version to be published; and both authors agree to be accountable for all aspects of the work in ensuring that questions related to the accuracy or integrity of any part of the work are appropriately investigated and resolved.

