## Supplemental figures, tables and protocol for "Charting Taxa in *Amanita* Section *Amidella* (*Basidiomycota : Amanitaceae*)"

Submitted to Mycological Progress

Authors:

1. Paulo Oliveira, ORCID 0000-0001-7979-6520
2. Ricardo Arraiano-Castilho, ORCID 0000-0001-8465-5909

Corresponding author: Paulo Oliveira, University of Évora, Biology Department, Évora, Portugal, Mediterranean Institute for Agriculture, Environment and Development (MED), Évora, Portugal, and Research Centre in Biodiversity and Genetic Resources (CIBIO), Vairão, Portugal. Email address:

Online Resource 1 – *Amidella* sequence database accessions (UDB refers to the UNITE database, otherwise except for one iNaturalist record, refer to GenBank; last search on December 2, 2024)

| Subsection | Stirp | Epithet | ITS | LSU | RPB2 | TEF1 | BTUB | etc. |
|---|---|---|---|---|---|---|---|---|
| Volvatae | Volvata | *clarisquamosa* | | AF024448 | | | | |
| | | aff. curtipes1 | KM052544 | KU139473 | | | | |
| | | *pseudovalens* | KJ950803 KJ950804 MW431348 MW431350 MW431352 MW431354 | KJ950782 KJ950783 MW431338 MW431339 MW431340 MW431341 MW431342 MW431343 MW431344 | | | | |
| | | *curtipes* | AY486235 AY486236 EF653963 KJ950805 MW494955 MW538529 PP680674 | EF653960 EF653961 KJ950784 MW494958 MW494959 | | | | |
| | | sp. QU06 | OQ946582 | | | | | |
| | | sp. whartonensis | ON129192 | MK277570 OQ754989 | | | | |
| | | *volvata* | MN722424 MT241886 MZ667952 MZ668151 OM809189 OP104057 OP104077/OP681759 OP541683 OP681749 OQ311333 | AF024485 OQ311333 | | | | |
| | Peckiana | *brunneomaculata* | MH508278 MH508279 MH508280 | MH486410 MH486411 MH486412 MH486413 | MH485892 MH485893 MH485894 MH485895 | MH508698 MH508699 MH508700 MH508701 | MH485428 MH485429 | |

| Subsection | Stirp | Epithet | ITS | LSU | RPB2 | TEF1 | BTUB | etc. |
|---|---|---|---|---|---|---|---|---|
| (Volvatae) | (Peckiana) | *lanigera* | MH508416 MH508417 MH508418 MH508419 MH508420 MH508421 ON794269 PP768065 | MH486617 MH486618 MH486619 MH486620 MH486621 MH486622 | MH486070 MH486071 MH486072 MH486073 MH486074 MH486075 | MH508876 MH508877 MH508878 MH508879 MH508880 | MH485590 MH485591 | |
| | | *peckiana* | MH281877 MK966422 OK316971 ON080993 OQ286517 OR026673 OR297618 PP436658 | HQ539720 MH620300 OQ286517 OR026673 | | | | |
| | Fulvisquamea | sp. 'Sarawak1' | | KY090936 KY090973 | | | | |
| | | *fulvisquamea* | HE814144 LC364238 OQ780689 OQ780690 OQ780691 OW732041 UDB01048294 | OQ780671 OQ780672 OQ780673 | OQ740051 OQ740052 OQ740053 | OQ740069 OQ740070 OQ740071 | OQ740087 | |
| Sagittariae | | aff. clarisquamosa2 | MK388161 | | | | | |
| | | aff. clarisquamosa1 | FJ375331 | | | | | |
| | | sp. OMS-013 | LC667043 | | | | | |
| | | sp. OMS-017 | LC667047 | | | | | |
| | | sp. montelongense | PP680673 | | | | | |
| | | aff. vaginata | OR978098 | | | | | |
| | | sp. sagittaria | MT073012 | MT073023 | | | | |
| | | sp. 17 GMB-2014 | KP012973 | KP012973 | | | | |
| Lepiotidae | Lepiotoides | *rufobrunnescens* | KT865208 KT865209 ON971238 ON971239 | KT865210 KT865211 OP363814 OP363815 | | | | |
| | | *parvirufobrunnescens* | | MW365377 | | | | |
| | | sp. JPN01/sp. OMS-015 | LC667045 MN794888/KT779088 MZ420670 | KT779075 MZ420670 | | | | |
| | | *lepiotoides* f. *subcylindrospora* | MN497357 MW431346 MW431347 | MN497358 MW431336 MW431337 | | | | |
| | | sp. QU05 | OQ946543 | OQ946543 | | | | |

| Subsection | Stirp | Epithet | ITS | LSU | RPB2 | TEF1 | BTUB | etc. |
|---|---|---|---|---|---|---|---|---|
| (Lepiotidae) | (Lepiotoides) | sp. IN25 | OM809223 | | | | | |
| | | sp. IN33 | MZ668207 | | | | | |
| | | sp. insolens | ON129236 iNAT:178300376 | OQ755010 | | | | |
| | | sp. texidella | | MT073025 | | | | |
| | | sp. canadensis01/sp. IN28 | MZ668090 OM972289 OR039770 | OR039770 | | | | |
| | | sp. 19 GMB-2014 | KP013009 | | | | | |
| | Avellaneosquamosa | aff. avellaneosquamosa2 | MH508258 | MH486379 | MH485873 | MH508681 | | |
| | | *parvicurta* | MH508489 MH508490 | MH486744 MH486745 | MH486168 MH486169 | | | |
| | | cf. *avellaneosquamosa*1 | MH508259 | AF024441 MH486380 | | MH508682 | | |
| | | aff. avellaneosquamosa3/ OMS-012 | KJ466418 LC667042 | KJ466483 | KJ466648 | KJ481982 | KJ466562 | |
| | | aff. avellaneosquamosa4/ aff. volvata3 | AB509467 JF723273 KJ466417 KM052554 | KJ466482 | KJ466647 KR824788 | KJ481981 | KJ466561 | |
| | | aff. volvata4 | KM052540 | | | | | |
| | | aff. volvata5 | KF245921 KM052547 | KF245905 | | | | |
| | | aff. avellaneosquamosa5 | MH508260 | | | | | |
| | | sp. IN31 | MZ262330 MZ668196 MZ668215 OR978102 PP436548 | | | | | |
| | | sp. S17/sp. IN41 | MK580700 MT241889 MT345230 MW464411 MZ668076 MZ668129 MZ668205 OM809210 OM972560 OP104059/OP681731 OP643234 OP681770 OP749011 OQ389389 OR800094 PP573989 | HQ539665 OQ754986 OQ755005 | | | | JF707921 HQ539773 HQ539876 HQ539980 |
| | | aff. rosea | | OQ628446 | | | | |
| | | sp. 12 GMB-2014 | KP012768 | | | | | |
| | Subviscosa | aff. subviscosa | MZ027793 | | | | | |

| Subsection | Stirp | Epithet | ITS | LSU | RPB2 | TEF1 | BTUB | etc. |
|---|---|---|---|---|---|---|---|---|
| (Lepiotidae) | (Subviscosa) | *subviscosa* s.l. (Benin) | HG995511 MW829797 MZ027794 MZ027795 MZ027796 MZ027797 MZ027798 MZ027799 MZ027800 MZ027801 UDB0800069 | HG995511 PQ528016 PQ528078 PQ528079 | | | | |
| | | *subviscosa* s.l. (Zambia) | FR731572 | | | | | |
| | | *subviscosa* s.l. (Zimbabwe) | UDB0800069 UDB0800100 | | | | | |
| | | aff. goossensiae KM 12 | | MT446289 | | | | |
| | | sp. TANZ01 | | MZ358873 | | | | |
| | | sp. soil subtropical China 32 | OW732032 | | | | | |
| | | sp. soil subtropical China 33 | OW732033 | | | | | |
| | | sp. clone d28 3Ama | JQ347157 | | | | | |
| | | sp. 'Sarawak2' | | KY090972 | | | | |
| | | sp. Thai04 | MZ420676 | MZ420676 | | | | |
| | | sp. TUF116503 | UDB025267 | | | | | |
| | | sp. clone Ama3 | AB854647 | | | | | |
| | | sp. 'Jalisco' | MH160011 | | | | | |
| | Pseudomutabilis | sp. 'Lupane' | UDB0800134 | | | | | |
| | | *pseudorufobrunnescens* | | MK116903 MK120005 | | | | |
| | | aff. volvata2 | AB509773 | | | | | |
| | | sp. 'California' | OR767846 PP407486 | | | | | |
| | | aff. mutabilis | MH212007 | MH620228 MH620308 MK277550 | | | | |
| | | aff. volvata1 | | AF024487 | | | | |

| Subsection | Stirp | Epithet | ITS | LSU | RPB2 | TEF1 | BTUB | etc. |
|---|---|---|---|---|---|---|---|---|
| Ponderosae | Ponderosa | *ponderosa* | AY486233 AY486234 EF653962 KJ950785 KJ950786 KJ950787 KJ950788 KJ950789 KJ950790 KJ950791 KJ950792 KJ950793 KJ950794 KJ950795 KJ950796 KJ950797 KJ950798 KJ950799 KJ950800 KJ950801 KJ950802 MW431345 OQ120624 UDB0787041 | EF653957 EF653958 EF653959 KJ950779 KJ950780 KJ950781 MW431335 | | | | |
| | | sp. whetstoneae | KC841903 KC841904 KC841905 KC841906 KC841907 KC855221 KP284288 KU186814 KU186815 KU186816 KU186817 KU186818 KU186819 KX018800 KX061517 KX061518 KX061519 MK580730 MK580806 MZ267772 MZ411567 MZ668198 MZ668202 MZ668211 ON209212 OP001377 OP035793 OP101157 OP101161 OP749303 OR824528 PP156349 PP516769 PP516840 | AF042608 KP284289 KU186811 KU186812 KU186813 KX018806 KX018807 KX018808 KX061530 KX061531 KX061532 KX061533 MK277595 MZ411549 OP001370 OP001371 OP001372 OP001373 OP001374 OP001375 OP035793 OQ754985 OQ755004 OQ755014 | | | | |
| | Claristriata | aff. avellaneosquamosa6 | AY436447 | | | | | |
| | | aff. volvata6/sp. OMS-010 | AB015681 KF245922 KF245923 LC667040 OR852537 | KF245906 KF245907 | | | | |

| Subsection | Stirp | Epithet | ITS | LSU | RPB2 | TEF1 | BTUB | etc. |
|---|---|---|---|---|---|---|---|---|
| (Ponderosae) | (Claristriata) | aff. volvata7 | KT779085 | KT779072 | | | | |
| | | aff. volvata8 | | KU139472 | | | | |
| | | *claristriata* | OQ780686 OQ780687 OQ780688 | OQ780668 OQ780669 OQ780670 | OQ740048 OQ740049 OQ740050 | OQ740066 OQ740067 OQ740068 | OQ740085 OQ740086 | |
| | | sp. neocaesariensis/ sp. IN26 | MT073016 MZ242065 MZ668089 MZ668091 OM972273 OR297617 | MT073024 | | | | |
| | | sp. S19/aff. volvata9 | FJ596795 FJ596796 MF161251 MZ363616 | AF097387 AF097388 MK277571 MZ363616 OQ754990 OQ755007 OR344379 | | | | |
| | | sp. T51 | MK580770 MZ265216 OP541661 | MZ265216 | | | | |
| | Pinophila | aff. albidostipes | MH508502 | | | | | |
| | | *pinophila* | MH508503 MH508504 MH508505 | MH486758 MH486759 MH486760 | | | | |
| Baakani | | aff. goossensiae | MZ345339 MZ345340 MZ345341 | | | | | |
| | | cf. goossensiae | MZ345342 MZ345343 MZ345344 | | | | | |
| | | sp. 'Madagascar' | UDB013409 | | | | | |
| | | sp. Arora-01-560 | | HQ539698 | | | | HQ539802 HQ539909 HQ540012 |
| | | sp. Sharp2816 | UDB025774 | | | | | |
| | | sp. 'Naletali' s.l. | UDB0800029 UDB01047904 | | | | | |

### Online Resource 2

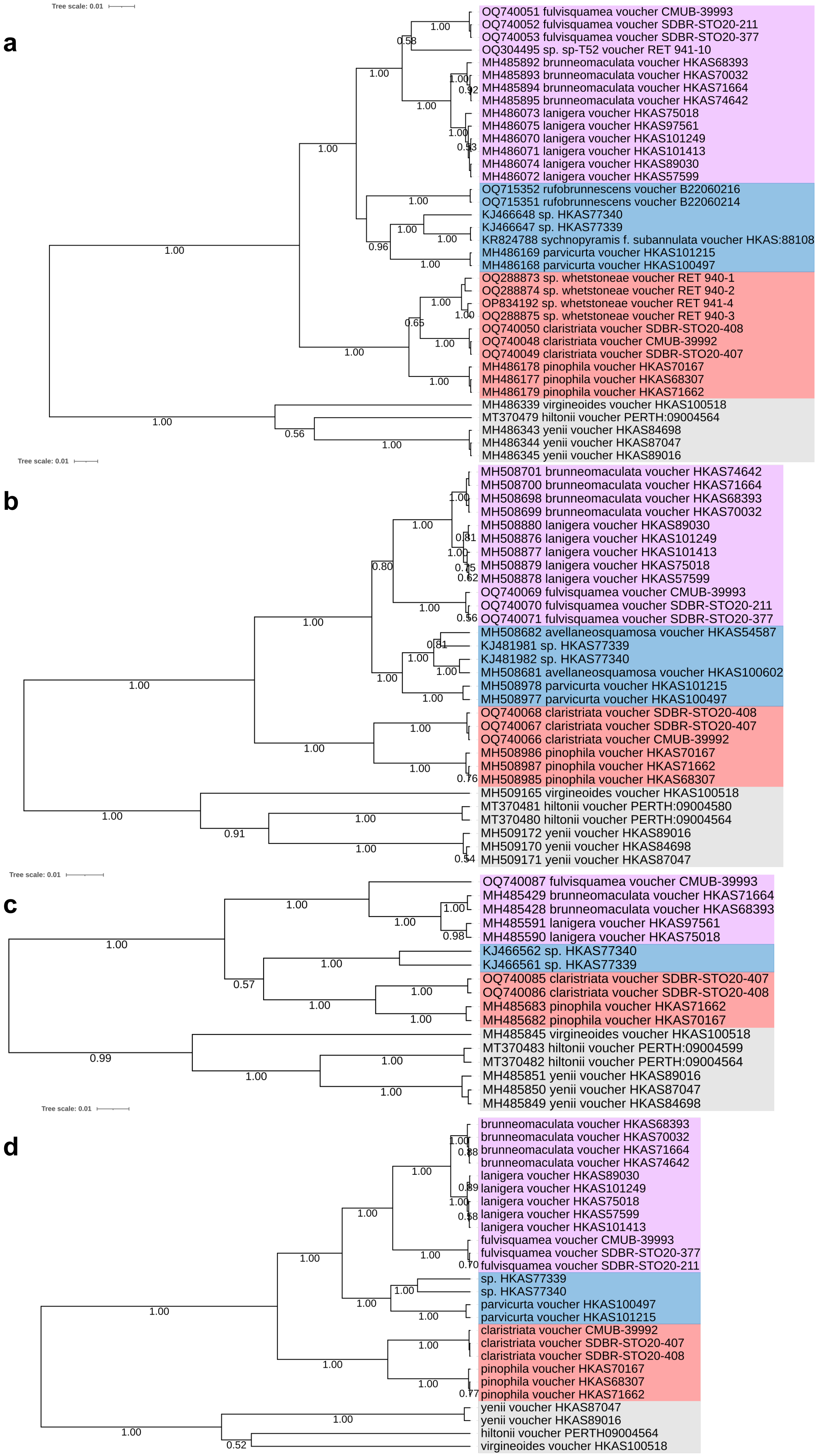

### Online Resource 3

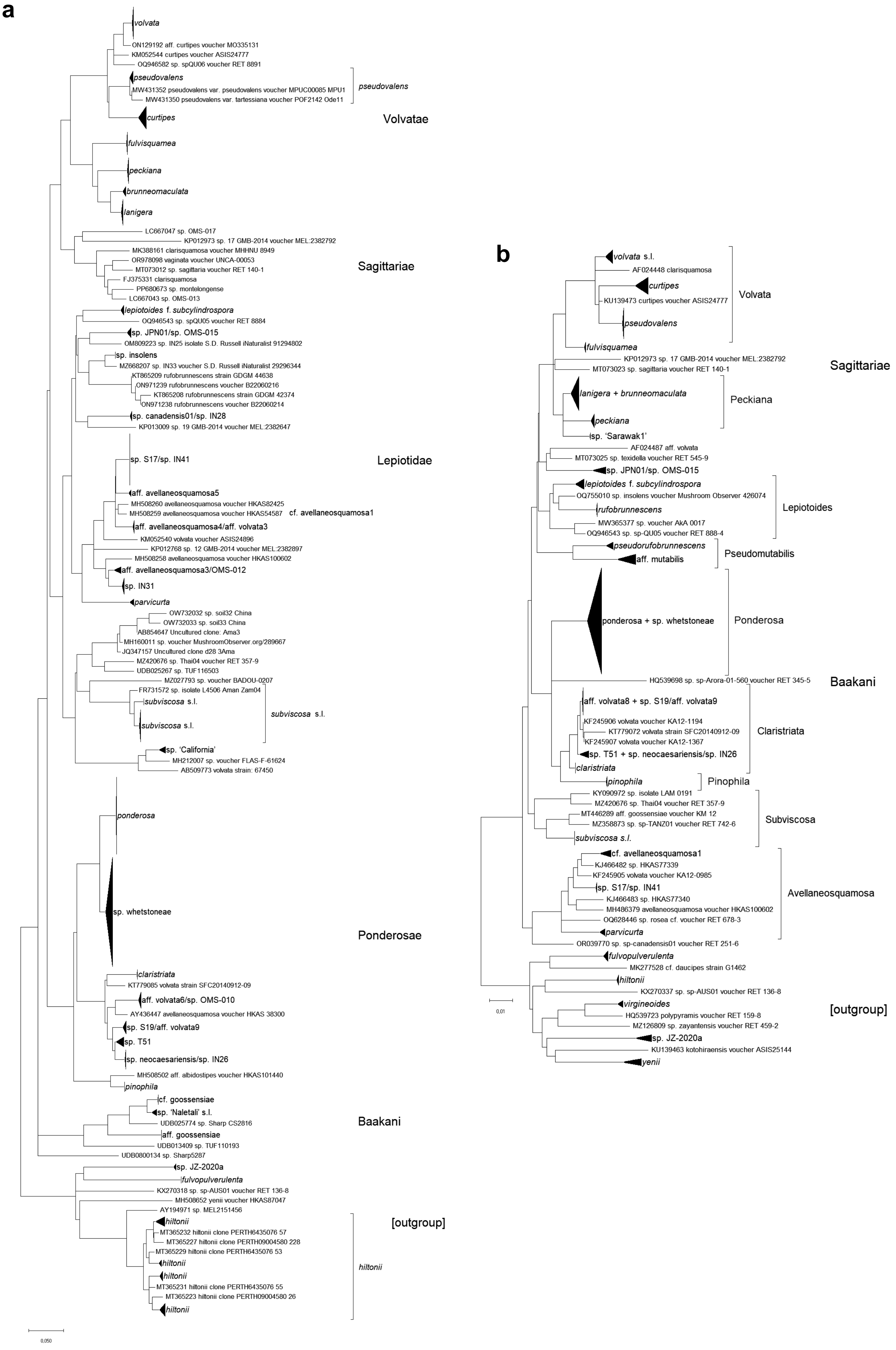

### Online Resource 4

**a**

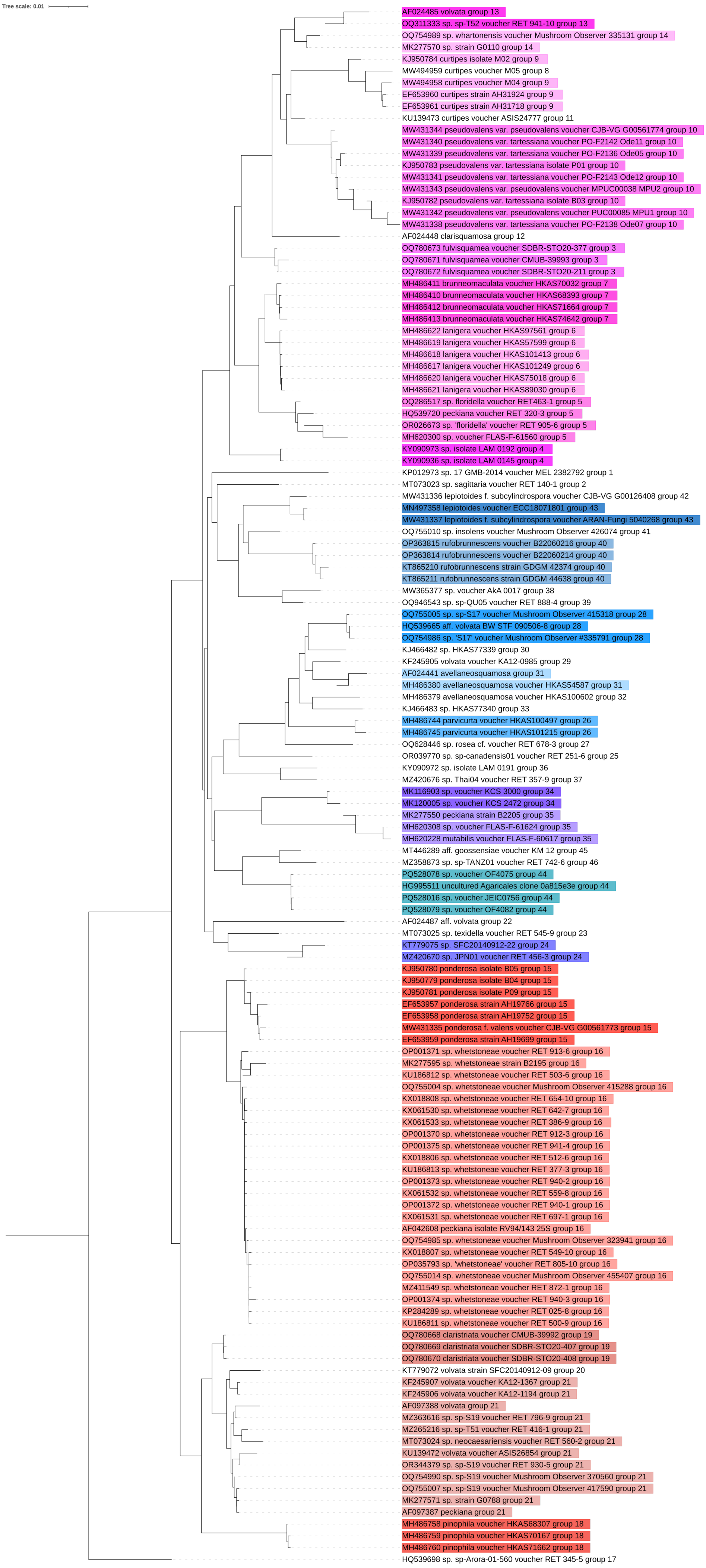

### Online Resource 4

**b**

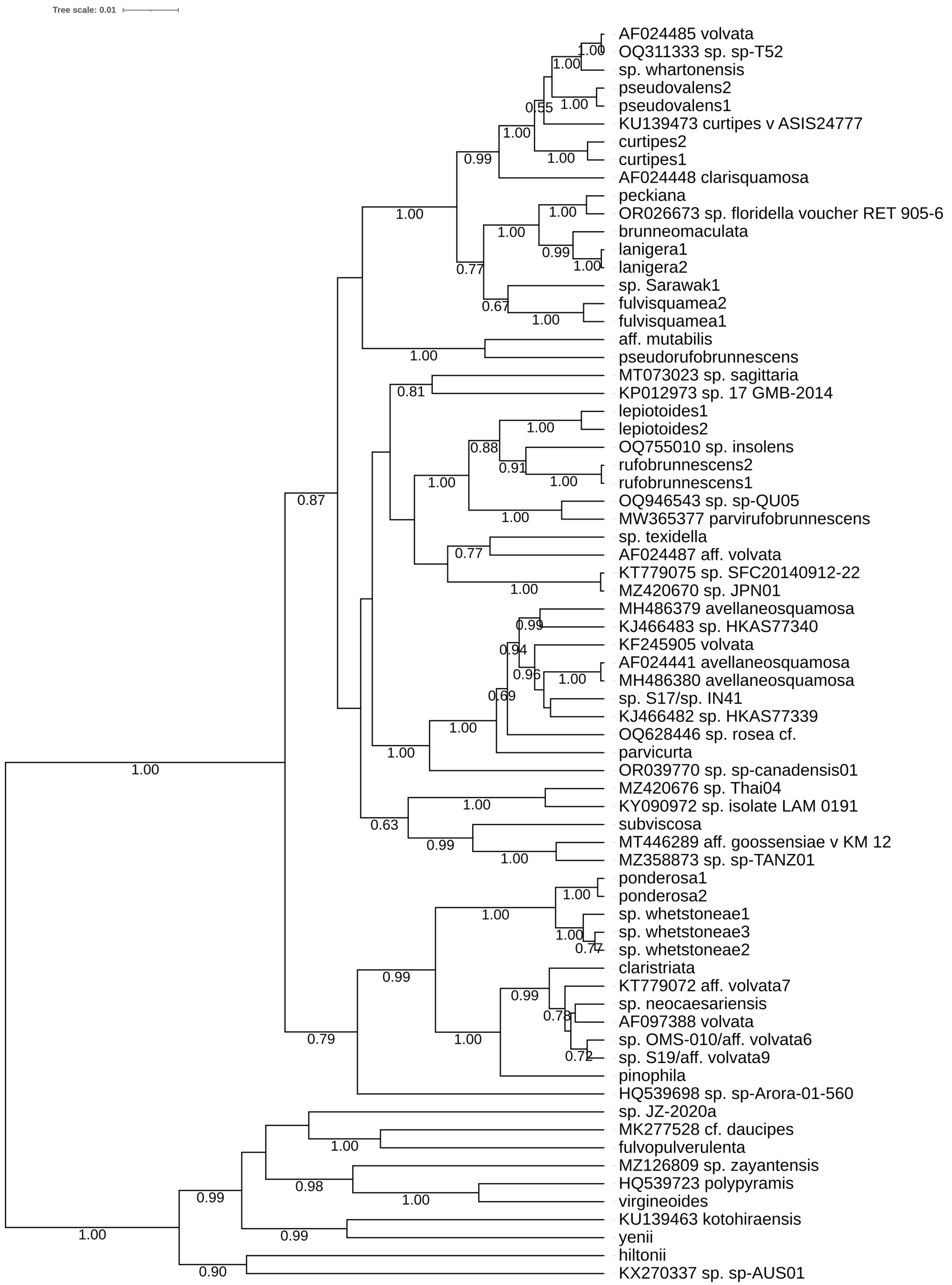

Online Resource 5 – Biogeography of the *Amidella* sequences, regions according to Loidi & Vynokurov (2024).

| Continent | Biogeographical regions (Kingdom) | Countries | Species clade | Subsection |
|---|---|---|---|---|
| Africa [a] | Guinean-Congolese (Paleotropical) | Cameroon | aff. goossensiae | Baakani |
| | | | cf. *goossensiae* | Baakani |
| | | | aff. goossensiae KM 12 | Lepiotidae |
| | Malagasy (Paleotropical) | Madagascar | sp. 'Madagascar' | Baakani |
| | Sudano-Zambezian (Paleotropical) | Benin | *subviscosa* s.l. | Lepiotidae |
| | | | aff. subviscosa | Lepiotidae |
| | | Tanzania | sp. TANZ01 | Lepiotidae |
| | | Zambia | *subviscosa* s.l. | Lepiotidae |
| | | | sp. Arora-01-560 | Baakani |
| | | Zimbabwe | sp. 'Lupane' | Lepiotidae |
| | | | *subviscosa* s.l. | Lepiotidae |
| | | | sp. naletali s.l. | Baakani |
| | | | sp. 'Sharp2816' | Baakani |
| Asia | Chinese-Japanese (Holarctic) | South Korea | aff. curtipes1 | Volvatae |
| | | | aff. avellaneosquamosa4/aff. volvata3 | Lepiotidae |
| | | | aff. volvata4 | Lepiotidae |
| | | | sp. JPN01/sp. OMS-015 | Lepiotidae |
| | | | aff. volvata5 | Lepiotidae |
| | | | aff. volvata8 | Ponderosae |
| | | | aff. volvata7 | Ponderosae |
| | | Japan/South Korea | sp. OMS-010/aff. volvata6 | Ponderosae |
| | | Japan | sp. OMS-013 | Sagittariae |
| | | | sp. OMS-017 | Sagittariae |
| | | | aff. volvata2 | Lepiotidae |
| | | Japan/China | aff. avellaneosquamosa3/OMS-012 | Lepiotidae |

| Continent | Biogeographical regions (Kingdom) | Countries | Species clade | Subsection |
|---|---|---|---|---|
| Asia | Chinese-Japanese (Holarctic) | China | *brunneomaculata* | Volvatae |
| | | | *clarisquamosa* | Volvatae |
| | | | *lanigera* | Volvatae |
| | | | aff. clarisquamosa1 | Sagittariae |
| | | | aff. clarisquamosa2 | Sagittariae |
| | | | aff. avellaneosquamosa2 | Lepiotidae |
| | | | sp. clone d28 3Ama | Lepiotidae |
| | | | aff. avellaneosquamosa5 | Lepiotidae |
| | | | *parvicurta* | Lepiotidae |
| | | | aff. volvata1 | Lepiotidae |
| | | | cf. *avellaneosquamosa*1 | Lepiotidae |
| | | | aff. avellaneosquamosa6 | Ponderosae |
| | | | *pinophila* | Ponderosae |
| | | | aff. albidostipes | Ponderosae |
| | | India | *parvirufobrunnescens* | Lepiotidae |
| | | | *pseudorufobrunnescens* | Lepiotidae |
| | Indo-Chinese (Paleotropical) | China | sp. soil subtropical China 32 | Lepiotidae |
| | | | sp. soil subtropical China 33 | Lepiotidae |
| | | | *rufobrunnescens* | Lepiotidae |
| | | Thailand | sp. TUF116503 | Lepiotidae |
| | | | sp. clone Ama3 | Lepiotidae |
| | | | sp. Thai04 | Lepiotidae |
| | | | *claristriata* | Ponderosae |
| | | Thailand (China, Japan) | *fulvisquamea* | Volvatae |
| | | Malaysia – Sarawak | sp. 'Sarawak1' | Volvatae |
| | | | sp. 'Sarawak2' | Lepiotidae |
| Europe | Eurosiberian (Holarctic) | Portugal | sp. montelongense | Sagittariae |
| | | Spain/France | *lepiotoides* f. *subcylindrospora* | Lepiotidae |
| | Eurosiberian/Mediterranean (Holarctic) | Portugal/France | *pseudovalens*[a] | Volvatae |

| Continent | Biogeographical regions (Kingdom) | Countries | Species clade | Subsection |
|---|---|---|---|---|
| (Europe) | Eurosiberian/Mediterranean (Holarctic) | Portugal/Spain/France/Italy (UK) | *ponderosa*[a] | Ponderosae |
| | Mediterranean (Holarctic) | Portugal/Spain | *curtipes*[a] | Volvatae |
| North America | Antillean-Mesoamerican (Neotropical) | Mexico | sp. 'Jalisco' | Lepiotidae |
| | Western North American (Holarctic) | USA | sp. 'California' | Lepiotidae |
| | North American Atlantic (Holarctic) | USA | *volvata* | Volvatae |
| | | | sp. whartonensis | Volvatae |
| | | | *peckiana* | Volvatae |
| | | | aff. vaginata | Sagittariae |
| | | | sp. sagittaria | Sagittariae |
| | | | aff. mutabilis | Lepiotidae |
| | | | sp. IN25 | Lepiotidae |
| | | | sp. canadensis01/sp. IN28 | Lepiotidae |
| | | | sp. IN31 | Lepiotidae |
| | | | sp. IN33 | Lepiotidae |
| | | | sp. S17/sp. IN41 | Lepiotidae |
| | | | sp. insolens | Lepiotidae |
| | | | sp. texidella | Lepiotidae |
| | | | sp. neocaesariensis/sp. IN26 | Ponderosae |
| | | | sp. S19/aff. volvata9 | Ponderosae |
| | | | sp. T51 | Ponderosae |
| | | | sp. whetstoneae | Ponderosae |
| | | Canada | sp. QU05 | Lepiotidae |
| | | | sp. QU06 | Volvatae |
| Oceania | Tasmanian-SE Australian (Australian) | Australia | aff. rosea | Lepiotidae |
| | North Australian (Australian) | | sp. 17 GMB-2014 | Sagittariae |
| | | | sp. 12 GMB-2014 | Lepiotidae |
| | | | sp. 19 GMB-2014 | Lepiotidae |

[a] Three species listed in Europe are known as fruitbodies in West Mediterranean Africa (Neville & Poumarat, 2004), Mediterranean region (Holarctic).

Online Resource 6 – List of described species tentatively in Section *Amidella*, currently (December 2024) without assigned DNA sequences.

| Continent | Countries | Name | Comment |
|---|---|---|---|
| Africa | Democratic Republic of Congo | *Amanita fulvosquamulosa* Beeli | |
| | | *Amanita subviscosa* Beeli | Sequences under this name from countries wide apart |
| | Madagascar | *Amanita irreperta* nom. prov. E.-J. Gilbert | Possibly the specimen classified as sp. 'Madagascar' |
| | Republic of the Congo | *Amanita floccosolivida* Beeli | possibly not *Amidella* |
| | | *Amanita olivacea* Beeli | |
| | Zambia | *Amanita subviscosa* sensu Pegler & Shah-Sm. | Basidiospores not fitting the description for the species (Tulloss & Yang 2024). One of the *subviscosa* s.l. is from Zambia |
| | "Congo", "Central Africa" | *Amanita goossensiae* Beeli | Sequences under that name in clades Lepiotidae and Baakani |
| Asia | Singapore | *Amanita duplex* Corner & Bas | |
| North America | Dominican Republic | *Amanita occidentalis* O. K. Mill. & D. J. Lodge | |
| | United States (Eastern) | *Amanita dolichopus* nom. prov. Tulloss | |
| | United States (Eastern), Canada | *Amanita pseudovolvata* nom. prov. Tulloss | |
| | United States (Western), Mexico | *Amanita fallax* nom. prov. Tulloss & G. Wright | Could be either sp. 'California' or (as suggested in Tulloss & Yang 2024) sp. 'Jalisco' |
| Oceania | Australia (NSW) | *Amanita pallidogrisea* A. E. Wood | possibly not *Amidella* |
| | Australia (Tasmania) | *Amanita grisea* Massee & Rodway | |
| | Australia (Victoria) | *Amanita curta* (Cooke & Massee) E.-J. Gilbert | |

### Online Resource 7

#### Volvatae

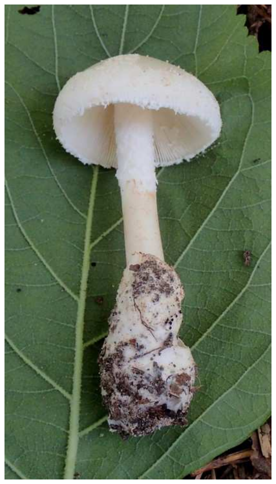

*volvata*
OQ311333

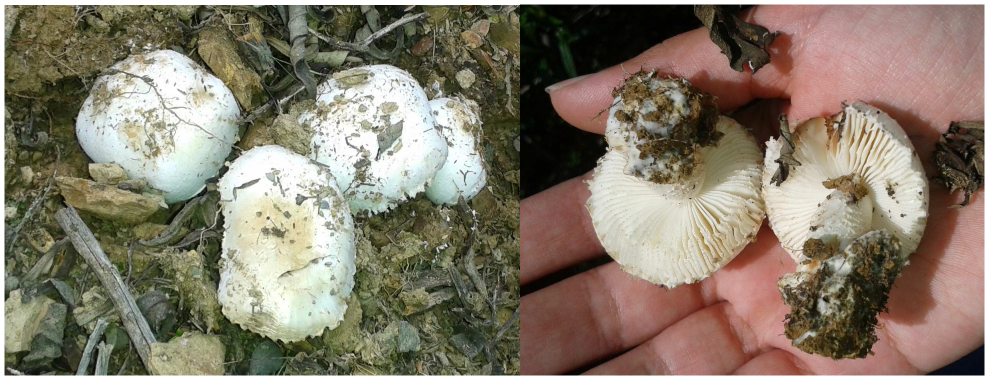

*pseudovalens* var. *tartessiana*
MW431341

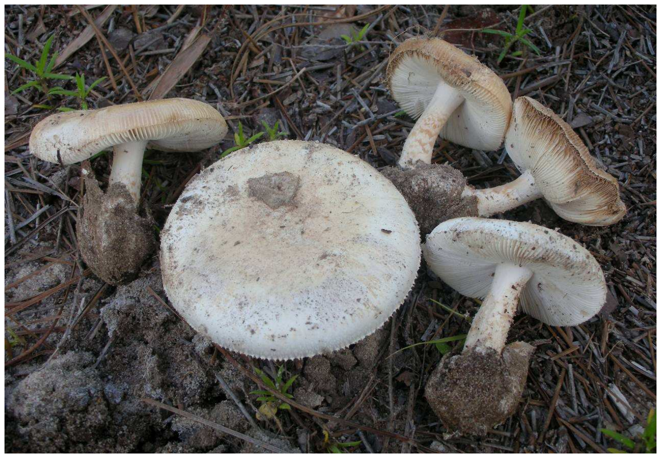

*peckiana*
OR026673

#### Sagittariae

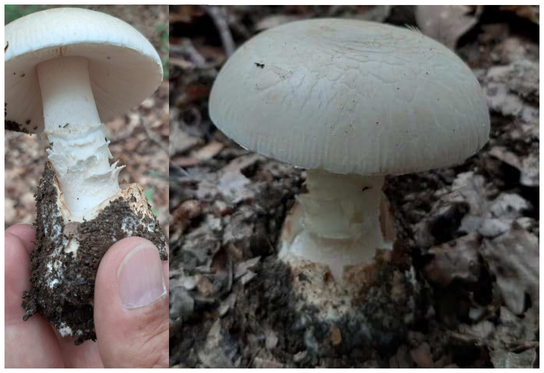

sp. montelongense
PP680673

#### Lepiotidae

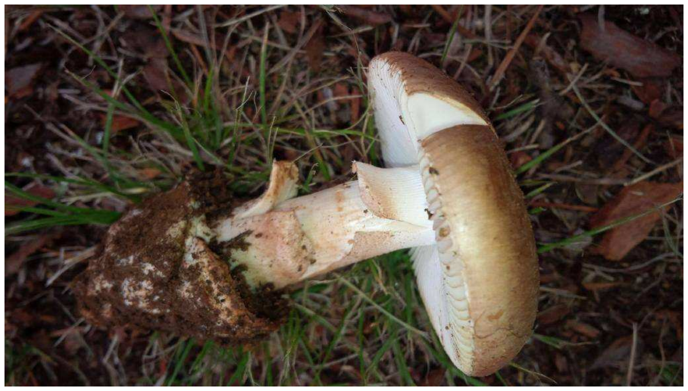

aff. rosea
OQ628446

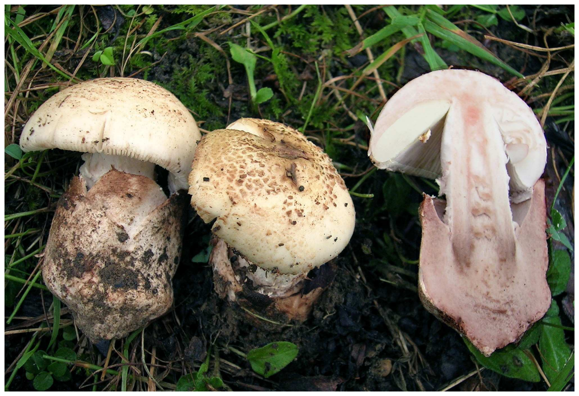

*lepiotoides* f. *subcylindrospora*
MW431337 MW431347

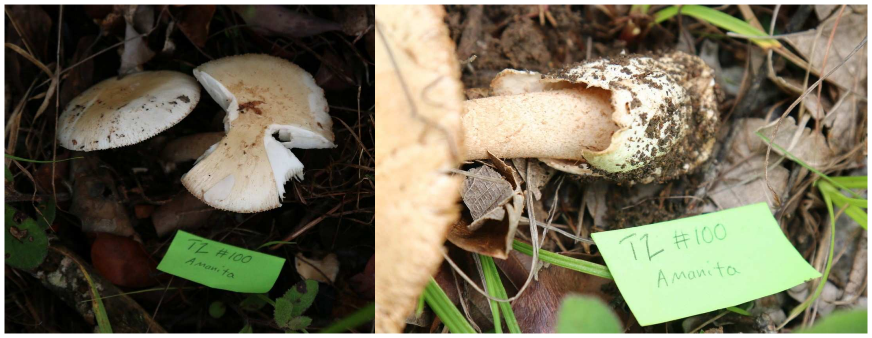


sp. TANZ01
MZ358873

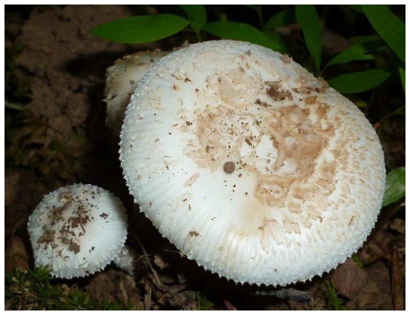

sp. S17/sp. IN41
OQ755005

#### Ponderosae

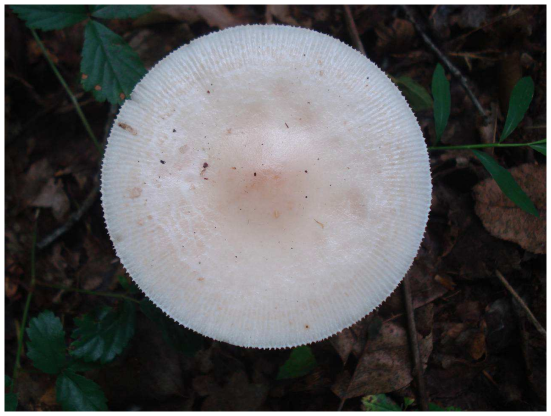

sp. whetstoneae
KX061533 KX061519

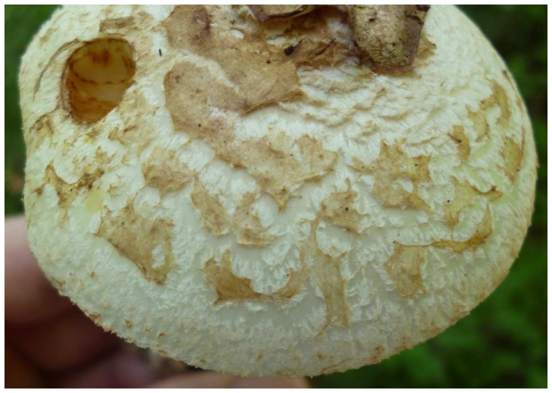

sp. S19/aff. volvata9
OQ754990

#### Photo credits

Note: the photos collected from Mushroom Observer, marked MO in the following list, are licensed through the Creative Commons BY-SA 3.0 https://creativecommons.org/licenses/by-sa/3.0/

*volvata*: MO 457058 Amanita sp. 'T52' (image 1341369) Photo by Ron Pastorino (2021) https://mushroomobserver.org/obs/457058
*pseudovalens* var. *tartessiana*: Amanita pseudovalens var. tartessiana. Photo by Ana C. Silva (2015), used with permission.
*peckiana*: MO 136750 Amanita floridella (image 338679) Photo by Ron Pastorino (2013) https://mushroomobserver.org/obs/136750
sp. montelongense: Amanita sp. PA-2024a. Photo by Helder Rodrigues (2022), used with permission.
aff. rosea: MO 197733 Amanita rosea (image 503402). Photo by Lucy Albertella (2015) https://mushroomobserver.org/obs/197733
*lepiotoides* f. *subcylindrospora*: Amanita lepiotoides f. subcylindrospora. Photo by Pedro Arrillaga (2006), used with permission.
sp. TANZ01: MO 240597 Amanita sp. 'TANZ01' (images 622776/9) Photos by Johannes Harnisch (2016) https://mushroomobserver.org/obs/240597
sp. S17/sp. IN41: MO 415318 Amanita sp. 'S17' (image 1199497) Photo by I. G. Safonov (2020) https://mushroomobserver.org/obs/415318
sp. whetstoneae: MO 175265 Amanita whetstoneae (image 448836) Photo by David Wasilewski (2014) https://mushroomobserver.org/175265
sp. S19/aff. volvata9: MO 370560 Amanita sp. 'sp-S19' (image 1048387) Photo by I. G. Safonov (2019). https://mushroomobserver.org/obs/370560

### Online Resource 8

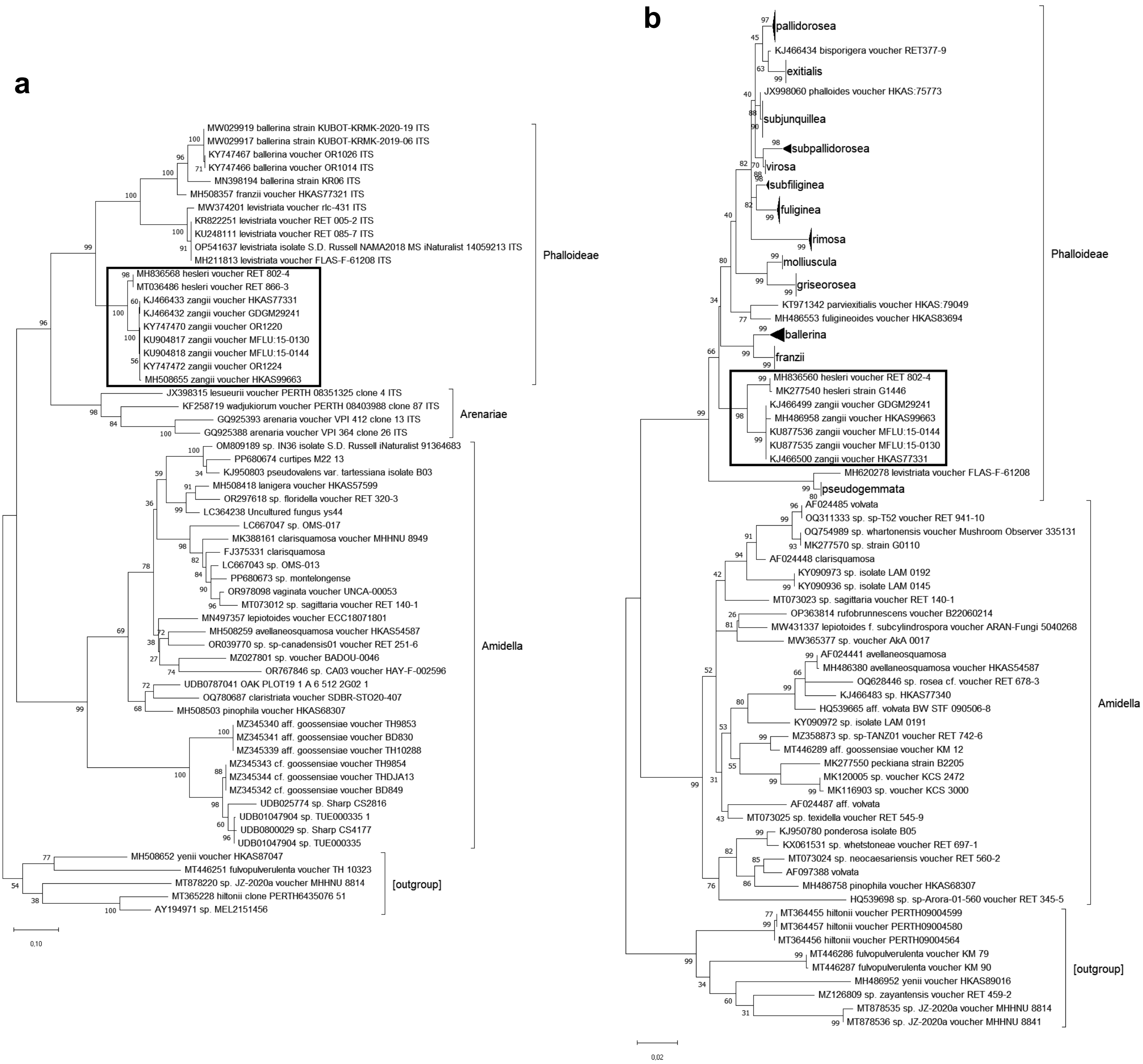

### Online Resource 9

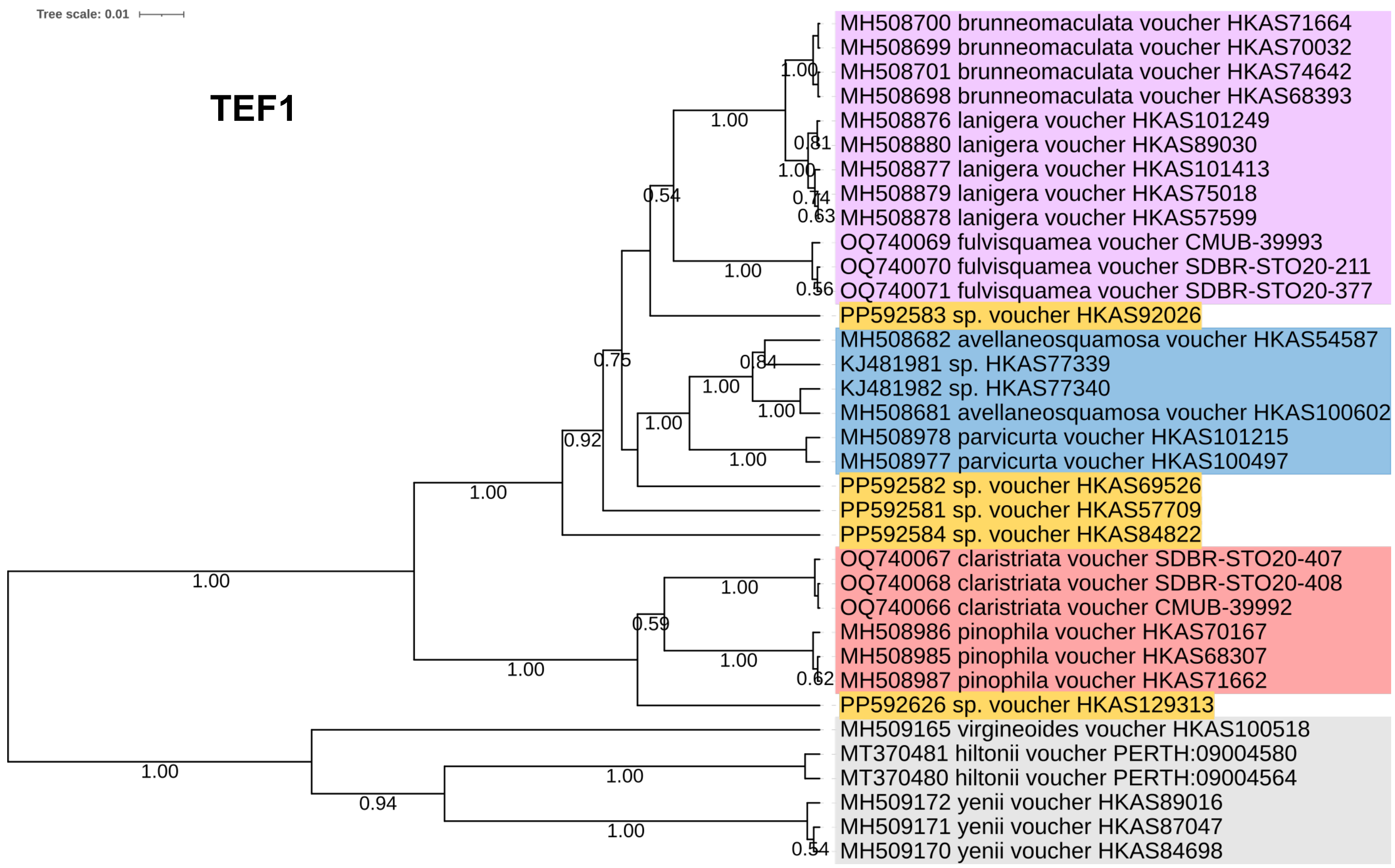

### Online Resource 10

Targeted PCR amplification from old specimens

#### Introduction

This procedure is expected to facilitate the retrieval of DNA sequences from old specimens where the degree of DNA degradation is such that genomic DNA fragments of enough length for successful amplification form a residual fraction.

It encompasses two rounds of amplification, the second one consisting of reamplification from samples of the previous round, using taxonomy-selective primers. The latter primers can be used already from the outset.

The choice of primers must be such that the resulting amplicons are relatively short (<400 bp).

A PCR block that can run at different temperatures simultaneously (temperature gradient blocks, for example) will facilitate optimising the primer annealing temperatures.

This procedure enabled the sequencing of ITS regions from specimens over 50 years old (Arraiano-Castilho et al., 2022).

#### Protocol

1. Prepare an enzyme mix (without primers). For the whole experiment, the $Mg^{2+}$ concentration in the final PCR reaction is set to 2.5 mM, adjusting the volume of $MgCl_2$ to be added depending on the composition of the enzyme buffer. The water volume is such that the enzyme mix fulfils 60% of the final volume. A standard Taq polymerase can be used.
2. Prepare the primer mixes, such that they form 30% of the total volume in each PCR reaction and the final concentration of each primer is 0.5 µM each.
3. First round: set up a 10 µL PCR reaction containing 1 µL of genomic DNA extract, 3 µL of primer mix, and 6 µL of the enzyme mix. Cover each reaction with 10 µL mineral oil.
    a. Ampflify over 40 cycles, allowing 1 minute for the elongation step in each cycle and a final elongation step of 5 minutes.
4. Second round: set up the PCR reactions with 3 µL water, 1 µL PCR solution from the corresponding first-round reaction, 12 µL primer mix, and 24 µL enzyme mix.
    a. Amplify over 30 cycles, allowing 15–30 seconds for the elongation step in each cycle and a final elongation step of 5 minutes.
5. Analyse the PCR reactions, using a 3 µL sample from each one, by agarose gel electrophoresis (at 1.25% in TBE 0.5X).

##### Example (possible experiment)

###### Materials and hypotheses

Two DNA extracts from dried vouchers belonging in *Amanita* section *Amidella*:

- Voucher #1 could be from clade Volvatae or clade Baakani (there are primers for the ITS2 region in both clades);
- Voucher #2 could be from clade Lepiotoides or clade Avellaneosquamosa (there are primers for the LSU gene only for the latter).

#### Strategies

For #1, amplify with universal primers in the first round, and then with taxon-specific primers in the second round (10 + 2x40 µL). Expected results: positive only for one of the tests (Baakani vs. Volvatae). A positive control such as the one devised for #2 can be added in the second round (10 + 3x40 µL).

For #2, amplify with an Avellaneosquamosa-specific primer in the whole experiment (test), using section-wide primers as positive controls (2x(10 + 40 µL)). Expected results: positive for the control, but positive for the test only if belonging to clade Avellaneosquamosa.

#### Setup

##### *Enzyme mix (120 µL):*

Enzyme buffer 10x (containing 15 mM $Mg^{2+}$) ......20 µL

$MgCl_2$, 25 mM........................................................8 µL

dNTP mix, 4 x 10 mM ............................................4 µL

Taq polymerase, 5 U/µL .......................................1 µL

Water..................................................................87 µL

Note: if a positive control is added for voucher #1, this mix will be scaled up to 150 µL.

##### *Primers:*

#### #1:

*First round (single reaction)*
forward ITS3 5’GCATCGATGAAGAACGCAGC, reverse ITS4 5’TCCTCCGCTTATTGATATGC

Mix: 0.5 µL each primer at 10 µM + 2 µL water.

*Second round (two reactions, one per test, possibly a third one for the positive control)*
Baakani

Forward 5’CAATYTTAAAATGCTTTYRG, reverse ITS4 5'TCCTCCGCTTATTGATATGC

Volvatae

Forward 5’AATGAATTAGCAGGGAWATATG, reverse ITS4 5'TCCTCCGCTTATTGATATGC

Each mix: 2 µL each primer at 10 µM + 8 µL water.

#### #2:

Test forward LR0R 5’ACCCGCTGAACTTAAGC, reverse 5’CTTYGAGAGCACACCACAAC

Control forward LR3R 5’GTCTTGAAACACGGACC, reverse 5’CAACATRCATGCTCTAC

Each mix (both rounds): 2.5 µL each primer at 10 µM + 10 µL water.

Online Resource 11

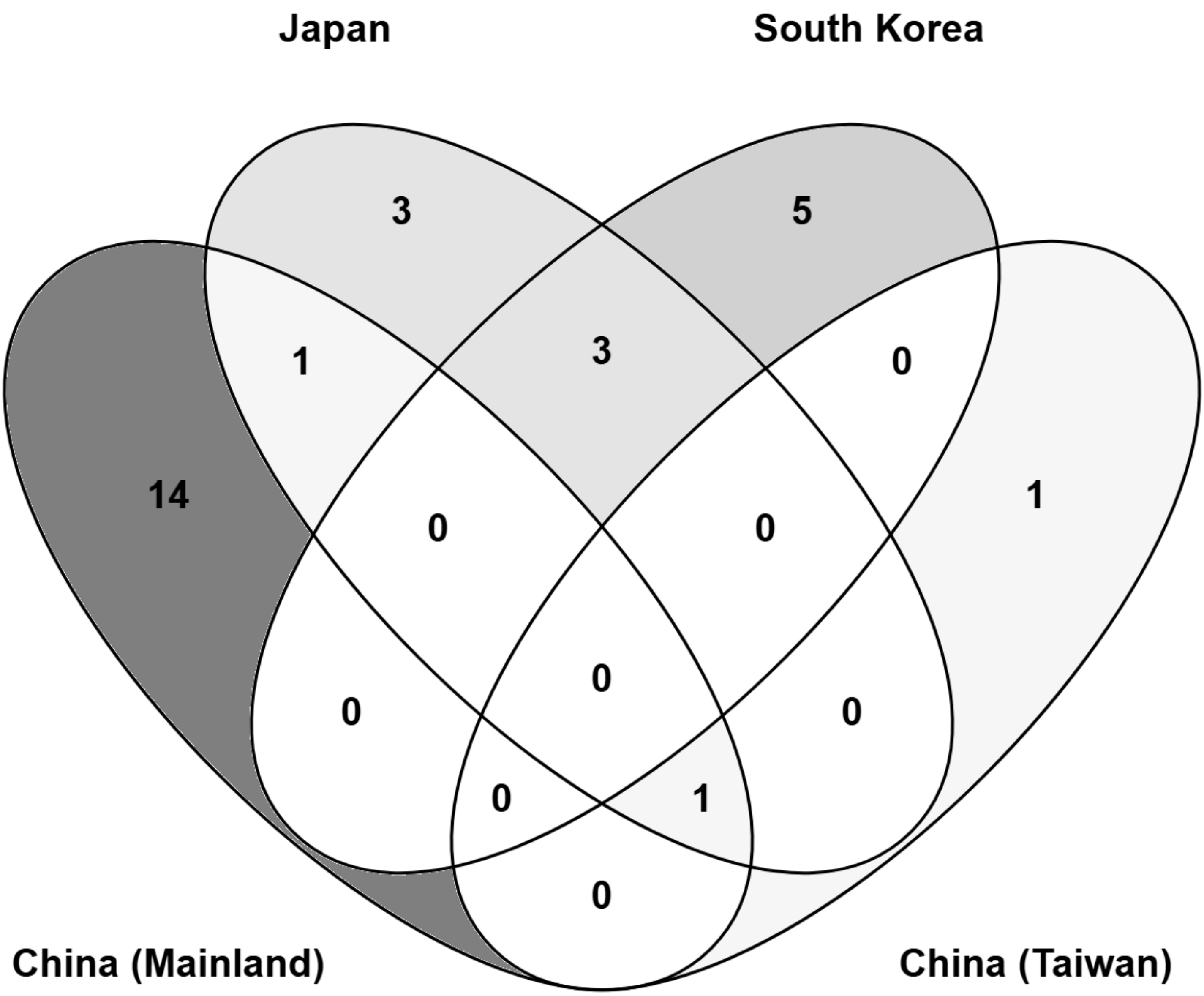

**Supplementary figures legends**

Online Resource 2. Phylogenetic trees obtained with BEAST, with posterior probabilities above 50% shown to the left of each node. The sequence labels have the GenBank accession numbers followed by unique identifiers, and the proposed clades are highlighted with the same background colours as in Figure 1. The scale bars are for substitutions per site. a) *RPB2* dataset, tree log likelihood –2,930.1. b) *TEF1* dataset, tree log likelihood –2,513.4. c) *BTUB* dataset, tree log likelihood –1,224.3. d) *LSU:RPB2:TEF1* concatenate dataset, tree log likelihood –7,681.8.

Online Resource 3. Phylogenetic trees obtained with MEGA (Kumar et al. 2024) using the Minimum Evolution Method with pairwise uncorrected p-distances, branches collapsed by sequence difference up to a common threshold and labelled according to the epithets listed in Table 4 and Online Resource 1. a) ITS1:ITS2 concatenate dataset, distance threshold 1.25%. b) ncLSU dataset, distance threshold 0.8%.

Online Resource 4. Phylogenetic trees for the ncLSU produced by clustering algorithms. a) ABGD (Puillandre et al. 2012) tree output, ingroup (*Amidella*) only, for the second init partition with MinSlope = 1.08, each sequence assigned to a group, with background colouring to highlight clusters of two or more sequences, leaving singletons uncoloured. Groups 9 and 21 are paraphyletic in this phylogeny. b) Phylogenetic tree obtained with BEAST, with posterior probabilities above 50% shown to the left of each node, from the aligned centroid sequences derived by USEARCH (Edgar 2010) from the ncLSU dataset, using the cluster_fast command (UCLUST algorithm as described in https://drive5.com/usearch/manual/uclust_algo.html) and id option set at 0.992 (degree of similarity), tree log likelihood –6,309.7.

Online Resource 7. Representative collections classified in section *Amidella*, identified according to Table 4, with the corresponding GenBank accession numbers, and arranged according to their subclade affinities. All photos untouched, except for cropping as required.

Online Resource 8. Evolutionary analyses using MEGA (Kumar et al. 2024) designed to clarify the phylogenetic positions of the Hesleri clade (boxed). The scale bars are for substitutions per site. a) ITS1:ITS2 concatenate dataset phylogeny comprising members of sections *Phalloideae*, *Arenariae*, *Amidella* and (outgroup) *Roanokenses*, using the Maximum Likelihood method and General Time Reversible model of nucleotide substitutions with Gamma distribution modelling of evolutionary rate differences among sites and invariant sites, log likelihood –6,075.34, showing node bootstrap supports above 50% next to the branches. b) ncLSU dataset phylogeny comprising members of sections *Phalloideae*, *Amidella* and (outgroup) *Roanokenses*, same method as in a), log likelihood –6,023.87.

Online Resource 9. Phylogenetic tree based on the *TEF1* dataset in Online Resource 2b, with five additional sequences (orange background), tree log likelihood –3,096.5.

Online Resource 11. Venn diagram built with Venny version 2.1.0 (Oliveros 2007-2015) showing the distribution of 28 *Amidella* species from East Asia represented in this study.

**References (Online Resources)**

Arraiano-Castilho R, Silva AC, Vila-Viçosa C, et al (2022) The *Amidella* clade in Europe (*Basidiomycota*: *Amanitaceae*): clarification of the contentious *Amanita valens* (E.-J.Gilbert) Bertault and the importance of taxon-specific pcr primers for identification. Cryptogam Mycol 43:139–157. https://doi.org/10.5252/cryptogamie-mycologie2022v43a6.

Edgar RC (2010) Search and clustering orders of magnitude faster than BLAST. Bioinformatics 26:2460–2461. https://doi.org/10.1093/bioinformatics/btq461

Loidi J, Vynokurov D (2024) The biogeographical kingdoms and regions of the world. Mediterr Bot 45:e92333. https://doi.org/10.5209/mbot.92333

Kumar S, Stecher G, Suleski M, et al (2024) MEGA12: Molecular Evolutionary Genetic Analysis Version 12 for adaptive and green computing. Mol Biol Evol 41:msae263. https://doi.org/10.1093/molbev/msae263

Neville P, Poumarat S (2004) *Amidella*. In: *Amaniteae: Amanita, Limacella et Torrendia*. Candusso, Alassio, pp 645–699

Oliveros JC (2007-2015) Venny. An interactive tool for comparing lists with Venn's diagrams. https://bioinfogp.cnb.csic.es/tools/venny/index.html

Puillandre N, Lambert A, Brouillet S, Achaz G (2012) ABGD, Automatic Barcode Gap Discovery for primary species delimitation. Mol Ecol 21:1864–1877. https://doi.org/10.1111/j.1365-294X.2011.05239.x

Suchard MA, Lemey P, Baele G, et al (2018) Bayesian phylogenetic and phylodynamic data integration using BEAST 1.10. Virus Evol 4:. https://doi.org/10.1093/ve/vey016
